# Comparative analyses of disease-linked missense mutations in the RNA exosome modeled in budding yeast reveal distinct functional consequences in translation

**DOI:** 10.1101/2023.10.18.562946

**Authors:** Maria C. Sterrett, Lauryn A. Cureton, Lauren N. Cohen, Ambro van Hoof, Sohail Khoshnevis, Milo B. Fasken, Anita H. Corbett, Homa Ghalei

**Affiliations:** Department of Biology, Atlanta, Georgia, USA; Biochemistry, Cell and Developmental Biology Graduate Program, Atlanta, Georgia, USA; Genetics and Molecular Biology Graduate Program, Emory University, Atlanta, Georgia, USA; Department of Biochemistry, Emory University School of Medicine, Atlanta, Georgia, USA; Department of Microbiology and Molecular Genetics, The University of Texas Health Science Center at Houston, Houston, Texas, USA

**Keywords:** RNA exosome, RNA exosomopathy, budding yeast disease model, translation dysregulation

## Abstract

The RNA exosome is a multi-subunit, evolutionarily conserved ribonuclease complex that is essential for processing, decay and surveillance of many cellular RNAs. Missense mutations in genes encoding the structural subunits of the RNA exosome complex cause a diverse range of diseases, collectively known as RNA exosomopathies, often involving neurological and developmental defects. The varied symptoms suggest that different mutations lead to distinct *in vivo* consequences. To investigate these functional consequences and distinguish whether they are unique to each RNA exosomopathy mutation, we generated a collection of *in vivo* models by introducing pathogenic missense mutations in orthologous *S. cerevisiae* genes. Comparative RNA-seq analysis assessing broad transcriptomic changes in each mutant model revealed that three yeast mutant models, *rrp4-G226D, rrp40-W195R* and *rrp46-L191H,* which model mutations in the genes encoding EXOSC2, EXOSC3 and EXOSC5, respectively, had the largest transcriptomic differences. While some transcriptomic changes, particularly in transcripts related to ribosome biogenesis, were shared among mutant models, each mutation also induced unique transcriptomic changes. Thus, our data suggests that while there are some shared consequences, there are also distinct differences in RNA exosome function by each variant. Assessment of ribosome biogenesis and translation defects in the three models revealed distinct differences in polysome profiles. Collectively, our results provide the first comparative analyses of RNA exosomopathy mutant models and suggest that different RNA exosome gene mutations result in *in vivo* consequences that are both unique and shared across each variant, providing further insight into the biology underlying each distinct pathology.

## INTRODUCTION

The steady-state levels of cellular RNAs are regulated through a delicate balance of transcription and decay. This balance is fine-tuned through co-and post-transcriptional events that include precise processing, decay, and quality control/surveillance of the RNA (Corbett 2018; Cramer 2019; Wolin and Maquat 2019). Beyond their impact on the transcriptome, the post-transcriptional regulatory events are critical to define the proteome in both time and space. The RNA exosome is an abundant, essential cellular machine that is a critical mediator of both RNA processing and decay. This ring-like, macromolecular complex is composed of nine structural subunits and a catalytic 3’-5’ exo-endoribonuclease (DIS3 or DIS3L in humans and Dis3/Rrp44 in budding yeast) (Mitchell et al. 1997; Makino et al. 2013). The subunits of the RNA exosome are highly conserved and were initially identified in *Saccharomyces cerevisiae* through a genetic screen for ribosomal RNA processing (*rrp*) mutants and biochemical analyses (Mitchell et al. 1996; Mitchell et al. 1997; Allmang et al. 1999b). The structural core of the RNA exosome is composed of three S1/KH cap subunits and a lower ring of six PH-like subunits. The 3-subunit cap is composed of EXOSC1/Csl4 (Human/*S. cerevisiae*), EXOSC2/Rrp4, and EXOSC3/Rrp40. The 6-subunit core is composed of EXOSC4/Rrp41, EXOSC5/Rrp46, EXOSC6/Mtr3, EXOSC7/Rrp42, EXOSC8/Rrp43, and EXOSC9/Rrp45. The structural cap and core subunits form a barrel-like structure through which RNA can be threaded in a 5’-3’ orientation. The DIS3, DIS3L, or Dis3/Rrp44 catalytic subunit is located at the bottom of the barrel and can process or degrade the RNA targets (**Figure 1A**). Structural studies of both yeast and human RNA exosome complexes have revealed conservation in the organization of the RNA exosome (**Figure S1A-B**) (Liu et al. 2006; Bonneau et al. 2009; Makino et al. 2013; Wasmuth et al. 2014; Zinder et al. 2016), beyond evolutionary sequence conservation of the subunits.

**Figure 1.**
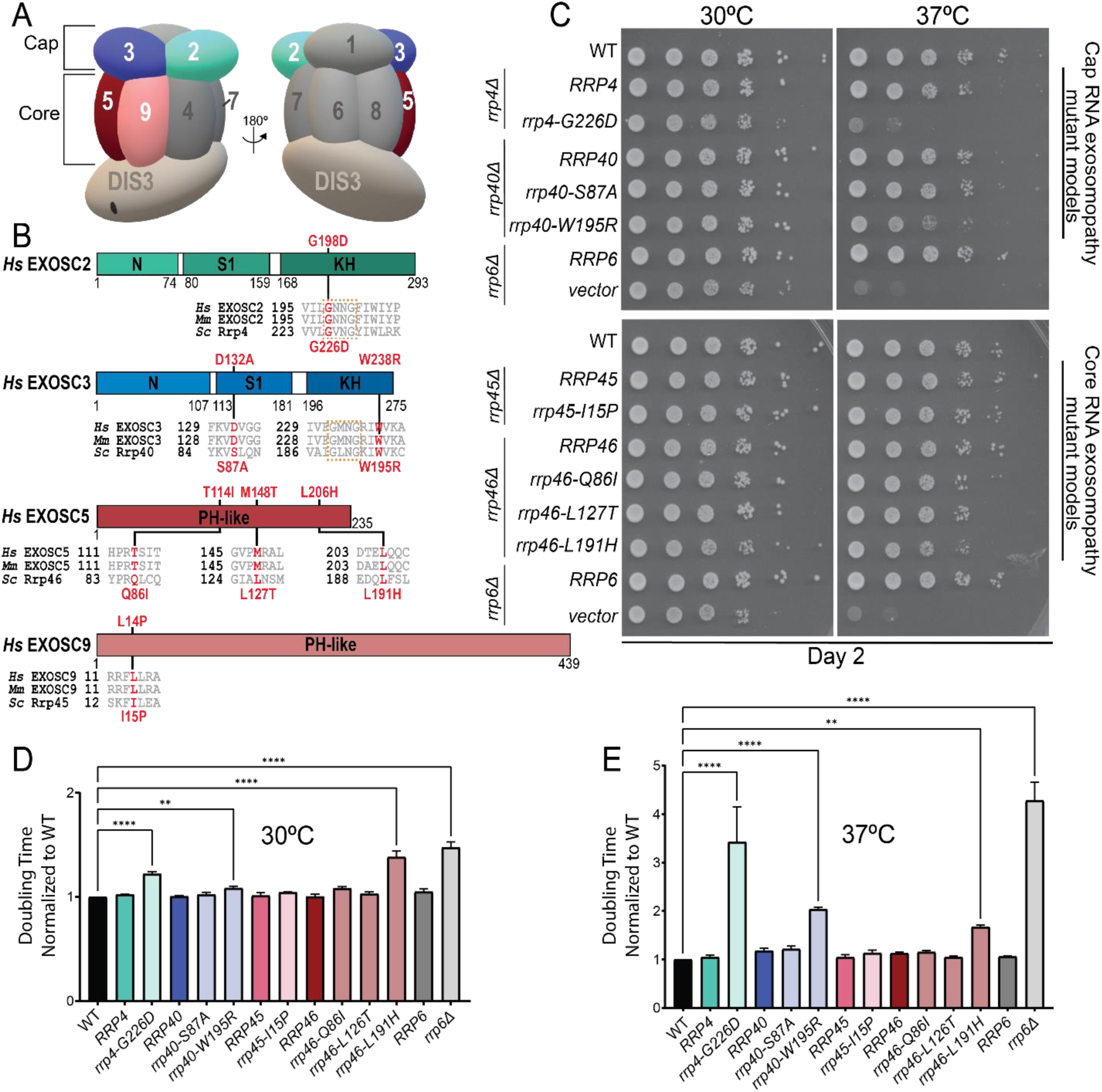
Overview of pathogenic amino acid substitutions in the human cap and core structural subunits of the RNA exosome. (**A**) Schematic view of the human RNA exosome with nine structural subunits (EXOSC1-9), denoted as 1-9, and one catalytic subunit (DIS3). (**B**) Domain maps of EXOSC2, EXOSC3, EXOSC5 and EXOSC9. EXOSC2 and EXOSC3, are composed of three domains: an N-terminal domain, a central putative RNA binding S1 domain, and a C-terminal putative RNA binding K homology (KH) domain. The “GxNG” motif in the KH domain of cap subunits is boxed in orange. EXOSC5 and EXOSC9 are composed of a singular PH-like domain. The positions of the RNA exosomopathy disease-linked amino acid substitutions in the human subunits are depicted above the domain structures in red. Sequence alignments of the orthologs from *Homo sapiens* (*Hs*), *Mus musculus* (*Mm*), and *S. cerevisiae* (*Sc*) reveal the high degree of conservation of the residues altered in disease (in red) and the sequences flanking these residues (in gray). The amino acid substitutions generated in the budding yeast Rrp orthologs for this study that correspond to the pathogenic amino acid substitutions are shown below the sequence alignments in red. The *rrp4-G226D*, *rrp40-W195R*, and *rrp46-L191H* mutant yeast cells, modeling *EXOSC2-G198D*, *EXOSC3-W238R*, and *EXOSC5-L206H* variants, respectively, exhibit (**C**) impaired growth in a solid media assay and (**D-E**) increased doubling times calculated from a liquid media assay at (**D**) 30°C and (**E**) 37°C. For these assays, the growth of RNA exosome deletion strains solely containing wild-type or mutant RNA exosome plasmid was analyzed by serial dilution and spotting of cells onto solid Leu^-^ media and in liquid media, at 30°C and 37°C. BY4741 cells were used as wild-type isogenic controls (labeled WT). The *rrp6Δ* deletion containing an empty vector or wild-type *RRP6* plasmid was used as a control. Growth measurements in liquid media used three biological replicates.

The RNA exosome plays a pivotal role in the processing, degradation, and surveillance of nearly every class of RNA in both the nucleus and cytoplasm (Schneider and Tollervey 2013; Kilchert et al. 2016; Morton et al. 2018). First discovered as a crucial complex required for proper maturation of ribosomal RNA (Mitchell et al. 1997; Allmang et al. 1999a), the RNA exosome has subsequently been shown to contribute to the processing of small nuclear RNAs (snRNAs), small nucleolar RNAs (snoRNAs) and transfer RNAs (tRNAs) (Allmang et al. 1999a; Chekanova et al. 2007; Gudipati et al. 2012; Schneider et al. 2012; Pefanis et al. 2014; Kilchert et al. 2016). In addition, the RNA exosome is critical for RNA homeostasis within the nucleus through targeting and degrading highly unstable species, such as cryptic unstable RNAs (CUTs) in *S. cerevisiae* and promoter upstream transcripts (PROMPTs) in human cells (Wyers et al. 2005; Preker et al. 2008; Kiss and Andrulis 2010; Parker 2012; Schneider et al. 2012; Belair et al. 2018). The RNA exosome also plays a crucial role in RNA surveillance in both the nucleus and cytoplasm, degrading aberrant RNAs (Belair et al. 2018). In addition to surveillance of misprocessed endogenous RNA species, the RNA exosome has been implicated in targeting foreign RNA through antiviral surveillance pathways (Molleston et al. 2016).

Though the RNA exosome is essential (Mitchell et al. 1997; Lorentzen et al. 2007; Hou et al. 2012; Lim et al. 2013; Pefanis et al. 2014; Brouze et al. 2025), recent clinical studies have identified pathogenic missense mutations in the structural subunit genes that result in distinct tissue-specific defects comprising a growing family of diseases termed RNA exosomopathies (Fasken et al. 2020). Pathogenic missense mutations have been identified in the cap subunit genes *EXOSC1/2/3* and core subunit genes *EXOSC4/5/8/9* (Wan et al. 2012; Biancheri et al. 2013; Boczonadi et al. 2014; Eggens et al. 2014; Di Donato et al. 2016; Burns et al. 2017; Schottmann et al. 2017; Burns et al. 2018; Bizzari et al. 2020; Slavotinek et al. 2020; Somashekar et al. 2021; Fasken et al. 2024). Missense mutations in the genes encoding the EXOSC1 and EXOSC3 cap subunits and EXOSC8 and EXOSC9 core subunits cause forms of pontocerebellar hypoplasia (PCH), a severe disease characterized by early onset atrophy of the pons and cerebellum (Wan et al. 2012; Biancheri et al. 2013; Eggens et al. 2014; Halevy et al. 2014; Burns et al. 2017; Schottmann et al. 2017; Burns et al. 2018; Bizzari et al. 2020; Sakamoto et al. 2021; Somashekar et al. 2021; Damseh et al. 2023). Missense mutations in the gene encoding the EXOSC5 core subunit are linked to a disease characterized by cerebellar atrophy, spinal muscular atrophy (SMA)-like motor delays and hypotonia (Slavotinek et al. 2020). In contrast to most of the other mutations that primarily cause neurological defects, missense mutations in the gene encoding the EXOSC2 cap subunit are linked to a novel syndrome termed SHRF (<u>s</u>hort stature, <u>h</u>earing loss, retinitis pigmentosa, and distinctive facies) (Di Donato et al. 2016). While diverse in their clinical manifestations, typically, RNA exosomopathy missense mutations result in single amino acid substitutions in conserved domains of the structural subunits of the RNA exosome.

Several recent studies have begun to investigate the molecular consequences of the different pathogenic amino acid substitutions that occur in exosomopathies (Summarized in **Table S1**)(Morton et al. 2018; Fasken et al. 2020). Expression levels of EXOSC3-G31A and EXOSC3-W238R variants in a mouse neuronal line were reduced compared to wild-type mouse EXOSC3, suggesting that these amino acid substitutions could affect the stability of the subunit (Fasken et al. 2017). Additionally, analyses of PCH patient fibroblasts and skeletal muscle cells homozygous for the *EXOSC9-L14P* mutations revealed that the variant protein levels are decreased compared to EXOSC9 levels in control samples, suggesting the EXOSC9-L14P substitution impacts the stability of the subunit (Burns et al. 2018). Similarly, analyses of the EXOSC8-S272T variant in myoblasts and fibroblasts showed that the steady-state EXOSC8 level is significantly decreased compared to EXOSC8 in wild-type control cells (Boczonadi et al. 2014). In addition, in patient fibroblasts with mutations in *EXOSC3* and *EXOSC8*, the EXOSC9 protein level was reduced, suggesting that reduced levels of one RNA exosome subunit can destabilize the RNA exosome complex (Burns et al. 2018). However, reconciling the diverse clinical pathologies seen in RNA exosomopathies cannot be solely explained by reductions in levels of individual essential subunits and/or the level of the RNA exosome complex. Thus, modeling these missense mutations and performing functional *in vivo* studies is critical to reveal the biology underlying RNA exosomopathy diseases.

Analyses of some of these RNA exosomopathy mutations in genetic model systems reveal distinct molecular and functional consequences resulting from the different pathogenic amino acid substitutions (Fasken et al. 2017; Gillespie et al. 2017; Morton et al. 2020; Slavotinek et al. 2020; Sterrett et al. 2021). These studies suggest that both complex integrity and interactions with known RNA exosome cofactors may be differentially impacted by specific RNA exosomopathy mutations (Fasken et al. 2017; Gillespie et al. 2017; Morton et al. 2020; Slavotinek et al. 2020). Nuclear RNA exosome cofactors have been extensively characterized in the budding yeast system and include the exonuclease Rrp6, its obligate binding partner Rrp47, the essential RNA helicase Mtr4, and Mpp6 (de la Cruz et al. 1998; LaCava et al. 2005; Schneider and Tollervey 2013; Zinder and Lima 2017). Structural studies of both the budding yeast and mammalian RNA exosome show conservation in the composite sites through which these cofactors interact with the complex and suggest similar roles *in vivo* across the eukaryotic systems (Schuch et al. 2014; Falk et al. 2017; Wasmuth et al. 2017; Schuller et al. 2018; Weick et al. 2018). Any alteration in the RNA exosome levels or key cofactor interactions resulting from these amino acid substitutions would ultimately have an impact on the ability of the complex to process, degrade, or survey RNA targets in a cell. Changes in RNA target levels could profoundly impact certain tissues if key RNA classes or specific RNAs are misprocessed, defective RNA accumulates, and/or RNA homeostasis is dysregulated. While previous studies of these RNA exosomopathy mutations provide valuable characterization *in vivo*, there has not yet been a direct comparison of the defects in RNA exosome function across multiple cap and core RNA exosomopathy mutant models. A comparative assessment of how these exosomopathy amino acid substitutions affect the ability of the RNA exosome to process, degrade, and survey aberrant RNAs *in vivo* is critical to further understand the molecular consequences underlying each distinct exosomopathy disease pathology.

Here, we take advantage of the budding yeast model system to explore and compare the functional and molecular consequences of a set of pathogenic amino acid substitutions within the RNA exosome. Given that the RNA exosome was initially identified and has been most extensively studied in *Saccharomyces cerevisiae* (Mitchell et al. 1997; Allmang et al. 1999b; Sloan et al. 2012) and the high conservation in overall complex structure between the human and budding yeast RNA exosomes (Makino et al. 2013; Makino et al. 2015; Kowalinski et al. 2016; Zinder et al. 2016; Wasmuth et al. 2017; Gerlach et al. 2018), a budding yeast system provides a robust platform to comparatively assess the *in vivo* consequences of exosomopathy mutations. In this study, we generated and analyzed *S. cerevisiae* models of the exosomopathy amino acid changes identified in *EXOSC2*, *EXOSC3*, *EXOSC5*, and *EXOSC9* genes by mutating the orthologous budding yeast genes *RRP4*, *RRP40*, *RRP46,* and *RRP45.* We analyzed yeast cell growth and employed an unbiased RNA-seq approach to explore the consequences of these missense mutations. From these approaches, we detect the greatest functional defects in three of our mutant models, *rrp4-G226D, rrp40-W195R,* and *rrp46-L191H*. Comparative analyses of transcriptomes across these three models revealed some shared changes, particularly in coding and non-coding transcripts required for rRNA processing and ribosome biogenesis, suggesting potential defects in translation. We also identified differentially expressed genes unique to each of the three mutant models, suggesting that while there are some shared consequences, there are also distinct differences in RNA exosome function. Assessment of ribosome biogenesis and translation defects in the three models revealed shared defects in rRNA processing but distinct differences in polysome profiles. Our results represent an unbiased approach to comparatively characterize the molecular defects in the function of the RNA exosome across a collection of RNA exosomopathy mutant models and suggest distinct translational defects may underlie the unique molecular pathology of RNA exosomopathies.

## RESULTS

### RNA exosomopathy mutations modeled in *Saccharomyces cerevisiae* cause different growth phenotypes

We used the budding yeast model to assess the effects of each RNA exosomopathy mutation. As shown in **Figure 1B**, the residues that are substituted in RNA exosome subunits of individuals with RNA exosomopathy often lie within evolutionarily conserved regions of the proteins, allowing the variant to be readily modeled in *S. cerevisiae.* The EXOSC2 amino acid substitution Gly198Asp (G198D) (Di Donato et al. 2016) and the EXOSC3 amino acid substitutions Asp132Ala (D132A) and Trp238Arg (W238R) (Rudnik-Schoneborn et al. 2013; Eggens et al. 2014; Schottmann et al. 2017) occur in highly conserved domains of both cap subunits in similar regions. EXOSC2-G198D and EXOSC3-W238R affect a conserved structural “GxNG” motif within the RNA binding KH domain. The RNA exosomopathy-linked missense mutations identified in the core subunit genes *EXOSC5* and *EXOSC9* also result in amino acid substitutions in conserved domains of each protein. Distinct mutations in the *EXOSC5* gene result in amino acid changes Thr114Ile (T114I), Met148Thr (M148T), and Leu206His (L206H), which are located throughout the PH-like domain of the EXOSC5 protein. The RNA exosomopathy mutation in the *EXOSC9* gene results in the amino acid change Leu14Pro (L14P) located near the N-terminus of the EXOSC9 protein. Structural analysis of each RNA exosomopathy amino acid substitution suggests that these changes could affect inter-subunit binding interfaces or the conformation of the subunits themselves (Liu et al. 2006; Fasken et al. 2017; Fasken et al. 2020; Slavotinek et al. 2020; Sterrett et al. 2021).

We modeled the RNA exosomopathy mutations found in *EXOSC2/3/5/9* in the corresponding *S. cerevisiae* genes *RRP4/40/46/45*. The SHRF-linked *EXOSC2-G198D* mutation is modeled by the *rrp4-G226D* yeast cells. The PCH-linked *EXOSC3-D132A, EXOSC3-W238R* and *EXOSC9-L14P* mutations are modeled by the *rrp40-S87A, rrp40-W195R* and *rrp45-I15P* yeast cells, respectively. The *EXOSC5* RNA exosomopathy mutations *EXOSC5-T114I, EXOSC5-M148T,* and *EXOSC5-L206H* are modeled by the *rrp46-Q86I, rrp46-L127T,* and *rrp46-L191H* yeast cells, respectively. We first examined the functional consequences of *rrp4/40/45/46* yeast mutants. RNA exosome deletion strains (*rrp4Δ, rrp40Δ, rrp45Δ* and *rrp46Δ)* solely containing the *RRP4/40/45/46* wild-type gene or *rrp4/40/45/45* mutant gene were analyzed in both solid and liquid media growth assays (**Figure 1C-E**). The parental wild-type budding yeast strain (BY4741; WT) was included as an isogenic control for the RNA exosome deletion strains.

Previous studies have shown that the *rrp4-G226D, rrp40-W195R* and *rrp46-L191H* yeast mutants are viable but exhibit growth defects compared to the corresponding wild-type control cells (Fasken et al. 2017; Gillespie et al. 2017; Slavotinek et al. 2020; Sterrett et al. 2021). Consistent with these results, the *rrp4-G226D, rrp40-W195R,* and *rrp46-L191H* mutant cells show impaired growth at 37°C, and *rrp4-G226D* and *rrp46-L191H* cells exhibit impaired growth at 30°C on solid media compared to corresponding control cells (**Figure 1C**). The *rrp40-S87A, rrp46-Q86I, rrp46-L127T*, and *rrp45-I15P* mutant cells show no growth defects compared to corresponding wild-type control cells (**Figure 1C**). As a comparative control for cells disrupted for RNA exosome function, we also examined the growth of *rrp6Δ* cells, which lack the RNA exosome cofactor Rrp6 and exhibit growth defects (Briggs et al. 1998). The Rrp6 exonuclease is non-essential in budding yeast, however, Rrp6 assists the RNA exosome in targeting and degradation of several key transcript RNAs (Briggs et al. 1998; Schuch et al. 2014; Wasmuth et al. 2014; Marin-Vicente et al. 2015). As expected, the *rrp6Δ* cells show extremely poor growth at 37°C compared to control cells (**Figure 1C**). In liquid media, the *rrp4-G226D, rrp40-W195R* and *rrp46-L191H* mutant cells display significantly increased doubling times compared to wild-type *RRP4/40/45* control cells at both 30°C and 37°C (**Figure 1D,E**). Notably, the *rrp46-L191H* cells have the longest doubling time at 30°C and the *rrp4-G226D* cells have the longest doubling time at 37°C, similar in intensity to that of *rrp6Δ* cells.

Overall, the comparative growth analyses suggests that RNA exosomopathy mutations that alter different subunits of the RNA exosome have varied functional consequences *in vivo,* with the most significant growth defects observed for *rrp4-G226D, rrp40-W195R* and *rrp46-L191H* mutants. These three modeled mutations have previously been shown to have varying impacts on the protein levels of the individual yeast RNA exosome subunits (Fasken et al. 2017; Slavotinek et al. 2020; Sterrett et al. 2021), with the Rrp40-W195R variant showing the most significant protein instability at both 30°C and 37°C (Fasken et al. 2017). Given the *rrp40-W195R* cells show the mildest growth defect compared to the *rrp4-G226D* and *rrp46-L191H* cells, these data suggest that the observed growth defects are not simply due to varying levels of loss of the essential subunits and subsequent loss of the complex. Thus, these *S. cerevisiae* models can be used to assess the molecular consequences that may arise in the processing and/or degradation of RNA from these RNA exosomopathy mutations.

### Broad transcriptomic changes observed in *rrp4-G226D*, *rrp40-W195R* and *rrp46-L191H* mutants

To perform an unbiased analysis of the molecular consequences of the modeled RNA exosomopathy pathogenic amino acid substitutions, we performed RNA-seq analysis on the *rrp4-G226D*, *rrp40-S87A*, *rrp40-W195R, rrp45-I15P, rrp46-Q86I*, *rrp46-L127T,* and *rrp46-L191H* mutants and the corresponding wild-type controls. Differential transcript expression analysis of each mutant compared to its corresponding wild-type control revealed that *rrp4-G226D, rrp40-W195R* and *rrp46-L191H* mutant cells exhibit a large number of differentially increased or decreased transcripts compared to the corresponding wild-type *RRP4/40/46* control cells (**Figure 2A**). In contrast, the *rrp40-S87A, rrp45-I15P*, *rrp46-Q86I* and *rrp46-L127T* transcriptomes show only minor changes (**Figure S2)**. Unbiased principal component analyses (PCA) of the RNA-seq data revealed reproducibility amongst the RNA-seq biological replicates and confirmed that the *rrp4-G226D*, *rrp40-W195R* and *rrp46-L191H* transcriptomes are distinct from those of their wild-type controls (**Figure S3**).

**Figure 2.**
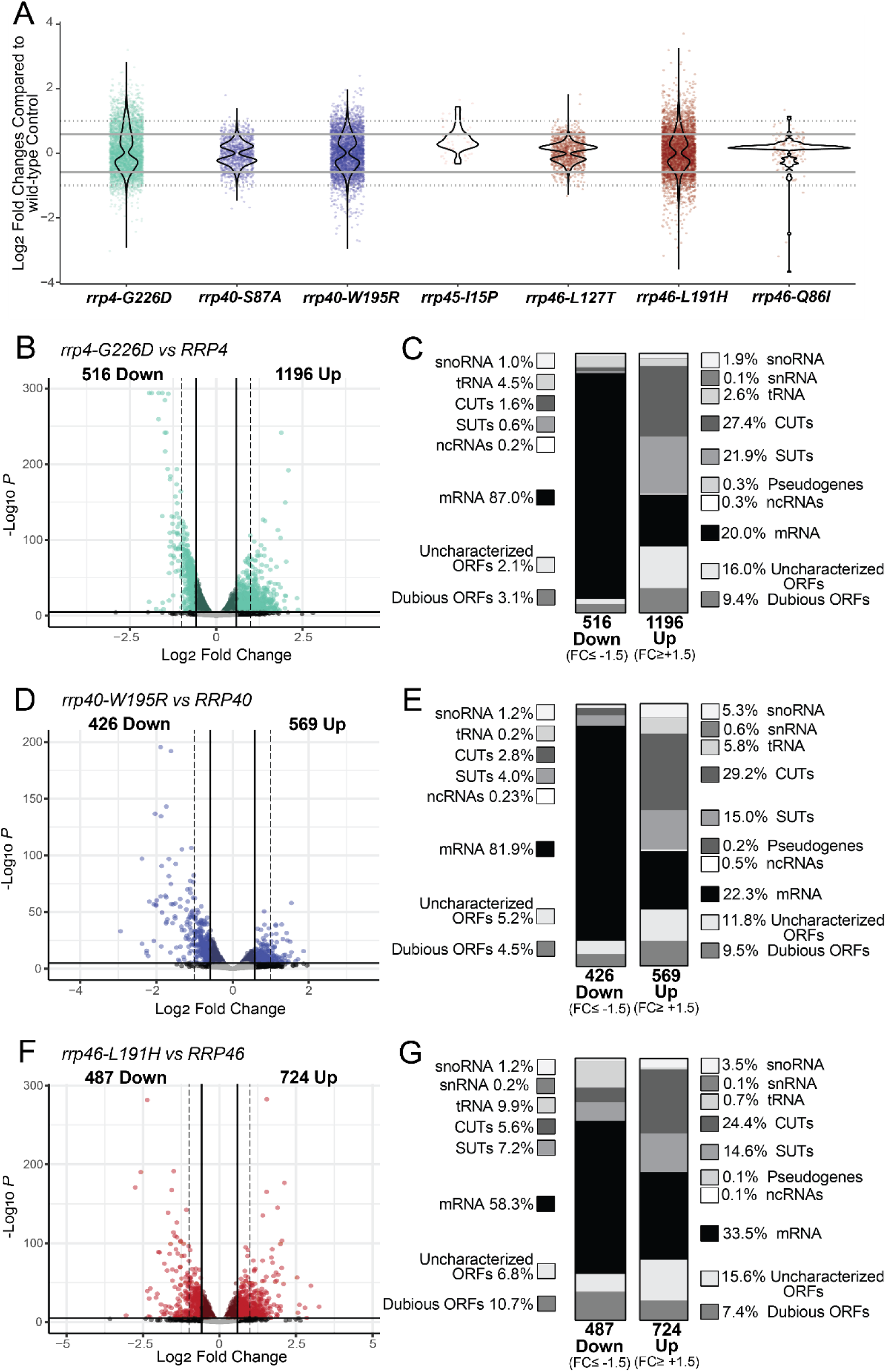
RNA-seq analysis of RNA exosomopathy mutant yeast models reveals large and distinct transcriptomic changes in the *rrp4-G226D, rrp40-W195R,* and *rrp46-L191H* mutant cells. (**A**) Violin plots showing the distribution of transcripts identified in differential analysis as significant (p<0.05) in each mutant compared to the corresponding wild-type control. The y-axis shows the Log2 Fold Change (LFC) for each transcript. The solid grey line demarcates a Fold Change of +1.5 or-1.5 (LFC=0.585 or-0.585). The dotted grey line marks a Fold Change of +2 or-2 (LFC=1 or-1). (**B, D, F**) Volcano plots of the differentially expressed transcripts and classification of RNA types in *rrp4-G226D, rrp40-W195R,* and *rrp46-L191H* mutant cells as labeled. (**C, E, G**) Stacked bar graph of the percentages of different RNA classes within the differentially expressed transcripts in *rrp4-G226D, rrp40-W195R,* and *rrp46-L191H* mutant cells as indicated. RNA classes are shown as percentages and include messenger RNAs (mRNA), small nuclear RNAs (snRNA), small nucleolar RNAs (snoRNA), transfer RNAs (tRNA), cryptic unstable transcripts (CUTs), stable unannotated transcripts (SUTs), other non-coding RNA (ncRNA; e.g., *TLC1*), pseudogenes, and uncharacterized or dubious open reading frames (ORFs). Vertical lines mark FC values of ±1.5 (straight line) and ±2 (dotted line).

RNA-seq analysis provides insight into the transcripts altered but does not provide any insight into which changes result directly from RNA exosome-mediated decay as compared to more systemic effects. Given the role of the RNA exosome in RNA decay, transcripts that show an increase in steady-state levels are more likely to be direct targets than transcripts that show a decrease in steady-state levels, so our analysis considers first the increased transcripts and then the decreased transcripts. Differential gene expression analysis of the *rrp4-G226D* cells reveals 1196 increased transcripts (FC ≥ 1.5, p<0.05) and 516 decreased transcripts (Fold change (FC) ≤-1.5, p<0.05) compared to control cells (**Figure 2B**). Increased transcripts are predominantly ncRNAs (**Figure 2C**), especially cryptic unstable transcripts (CUTs) and stable unannotated transcripts (SUTs), and the two most significantly increased mRNAs include *PIR3*, encoding a protein required for cell wall stability, and *DDR2*, encoding a multi-stress response protein (**Figure 2C**). In contrast, nearly 90% of decreased transcripts are mRNAs (**Figure 2C**), and the most significantly decreased transcripts include *SSA1, SSA2*, *HSC82*, and *HSP60,* encoding chaperones, as well as *RPS3* and *RPL15A,* encoding ribosomal proteins. In *rrp40-W195R* cells, 569 increased (FC ≥ 1.5, p<0.05) and 426 decreased (FC ≤-1.5, p<0.05) transcripts were identified (**Figure 2D**). Increased transcripts in *rrp40-W195R* cells compared to control (**Figure 2E**) are mostly ncRNAs, such as CUTs (29%), SUTs (15%), snoRNAs (5%), and tRNAs (6%). In contrast, over 80% of the decreased transcripts are mRNAs (**Figure 2E**), including *HIS4, URA1*, *URA4*, and *MDH2,* involved in metabolism, as well as *RPS13* and *RPS7B*, encoding ribosomal proteins. In *rrp46-L191H* cells, 724 increased transcripts (FC ≥ 1.5, p<0.05) and 487 decreased transcripts (FC ≤-1.5, p<0.05) were identified (**Figure 2F**). The increased transcripts in *rrp46-L191H* cells also include CUTs (30%), SUTs (15%), and mRNAs (30%) (**Figure 2G**), with *RPL18B,* encoding a ribosomal protein, and *CBT1*, encoding a protein involved in 5’ RNA end processing of mitochondrial cytochrome *b* mRNA that is linked to processing of rRNA, among the most significantly increased mRNAs. Approximately 60% of decreased transcripts in *rrp46-L191H* cells are mRNAs (**Figure 2G**), including *HIS4*, *URA1*, *URA4*, *BIO3*, *BIO4*, and *RIB4,* involved in metabolism, as well as *RPS3* and *RPL15A* mRNAs, encoding ribosomal proteins.

The overall comparison across the *rrp4-G226D, rrp40-W195R* and *rrp46-L191H* mutant cells reveals distinct changes in RNA categories affected (**Figures 2C**, **2E**, and **2G**), particularly in the distribution of increased ncRNAs and decreased mRNAs across these models. All three models show increases in CUTs and SUTs. As CUTs and SUTs are stabilized in RNA exosome mutants and crosslink to the RNA exosome (Wyers et al. 2005), these transcripts are likely direct targets of the RNA exosome, indicative of defective complex function. To compare changes in CUTs and SUTs across the three mutant models, we generated a heatmap of normalized fragments per Kilobase of transcript per Million mapped (FPKM) expression estimates (**Figure 3**). We included FPKM estimates of the *rrp6Δ* deletion strain as CUTs and SUTs were first identified in cells deleted for *RRP6* and Rrp6 activity is important for degradation of the CUTs (Wyers et al. 2005; Davis and Ares 2006; Xu et al. 2009). The *rrp6Δ* cells show a broad, indiscriminate increase in all CUTs and SUTs (**Figure 3**). The *rrp4-G226D* cells show a broad increase in CUTs and SUTs, while *rrp40-W195R* and *rrp46-L191H* cells also exhibit increases, but to a lesser extent than *rrp4-G226D* (**Figure 3**). Interestingly, not all the same CUTs and SUTs are changed in *rrp4-G226D*, *rrp40-W195R*, and *rrp46-L191H* cells. The *rrp4-G226D* cells exhibit the broadest increase in CUTs and SUTs among the three mutants, though certain groups remain unaffected. Similarly, *rrp40-W195R* and *rrp46-L191H* cells show both shared and unique changes in CUTs and SUTs (**Figure 3**).

**Figure 3.**
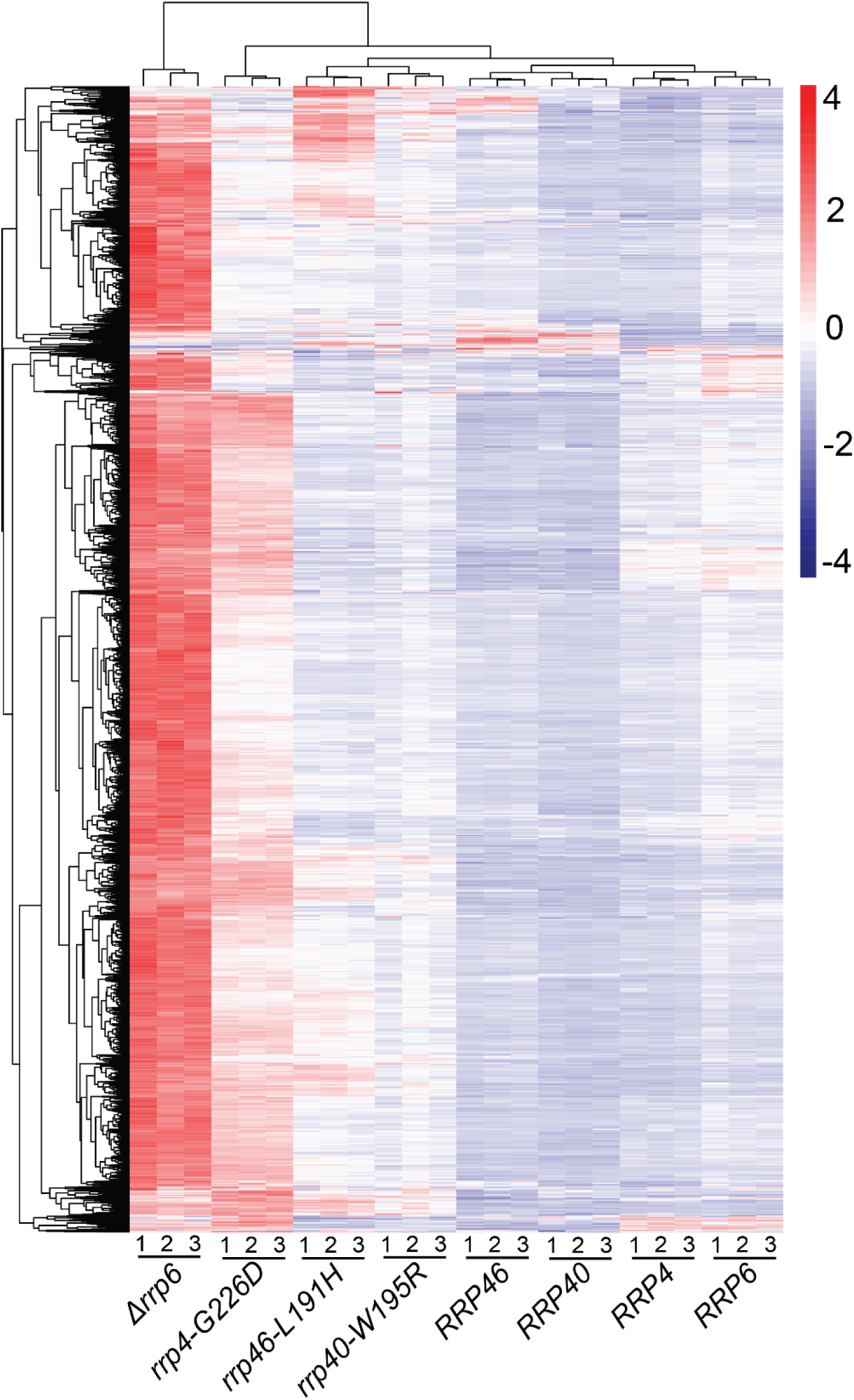
Heatmaps of RNA exosomopathy yeast mutants reveal an accumulation in CUTs/SUTs. Transcript level estimates of all cryptic unstable transcripts (CUTs) and stable unannotated transcripts (SUTs) are scaled for heatmap visualization. Coloring is a gradient of higher (red) to lower (blue) scaled values.

While most decreased transcripts in all three mutant models are mRNAs, the degree by which mRNAs, among other transcripts, are affected varies in each mutant. In *rrp46-L191H* cells, expressing a core subunit variant, ∼60% of the significantly reduced transcripts are mRNAs, compared to 80-90% in the *rrp4-G226D* and *rrp40-W195R* cells, expressing cap subunit variants. Additionally, in *rrp46-L191H* cells,

∼20% of total significantly decreased transcripts are tRNAs, CUTs, and SUTs, while these transcripts are significantly less affected in the other two mutants. These differences highlight the differential impact of cap vs core RNA exosome variants on the effected transcript types. These observations suggest that within the *rrp4-G226D, rrp40-W195R* and *rrp46-L191H* cells, RNA exosome targeting and degradation is impacted, yet in distinct ways, as all the direct RNA targets are not indiscriminately elevated across all three mutant models. These data strongly suggest that the broader differences in transcriptome profiles seen in these models (**Figures 2,3**) are unlikely to merely reflect the variable penetrance of the variants.

In summary, the diverse differentially increased/decreased transcripts observed across *rrp4-G226D, rrp40-W195R*, and *rrp46-L191H* cells highlight the distinct changes resulting from each RNA exosome variant, suggesting differential functional impacts of the variants *in vivo*.

### Comparative assessment of differentially expressed transcripts within *rrp4-G226D, rrp40-W195R* and *rrp46-L191H* mutants suggests shared impacts on metabolic pathways and rRNA modification and processing

To compare the molecular impacts of the modeled RNA exosomopathy mutations, we analyzed shared increased and decreased transcripts across the three models (*rrp4-G226D, rrp40-W195R* and *rrp46-L191H*) using UpSet plots (**Figure 4**). We identified 209 significantly increased transcripts (FC ≥ +1.5, p<0.05) across all three mutant models (**Figure 4A**), mostly CUTs and SUTs (**Figure 4B**), consistent with the trend observed for each individual mutant model. GO analysis revealed significant enrichment in rRNA modification (GO:0000154) (**Figure 4C**), possibly driven by the increased snoRNAs, which are required for rRNA modifications. We note that the total RNA-seq approach used here is not ideal for capturing all snoRNA changes in an unbiased manner as that was not the goal of this study and we did not employ a thermostable reverse transcriptase to accurately capture highly structured RNAs like snoRNAs (Boivin et al. 2018). Nonetheless, we observe a fraction of significantly differentially expressed snoRNA transcripts which are known to be direct targets of the RNA exosome (Webster and Ghalei 2023). Overall, the significant enrichment among the three mutant models in rRNA modification and processing (GO:0000154; GO:00031167; GO:0006364) (**Figure 4C**) strongly suggests significant impacts on ribosome biogenesis within these models.

**Figure 4.**
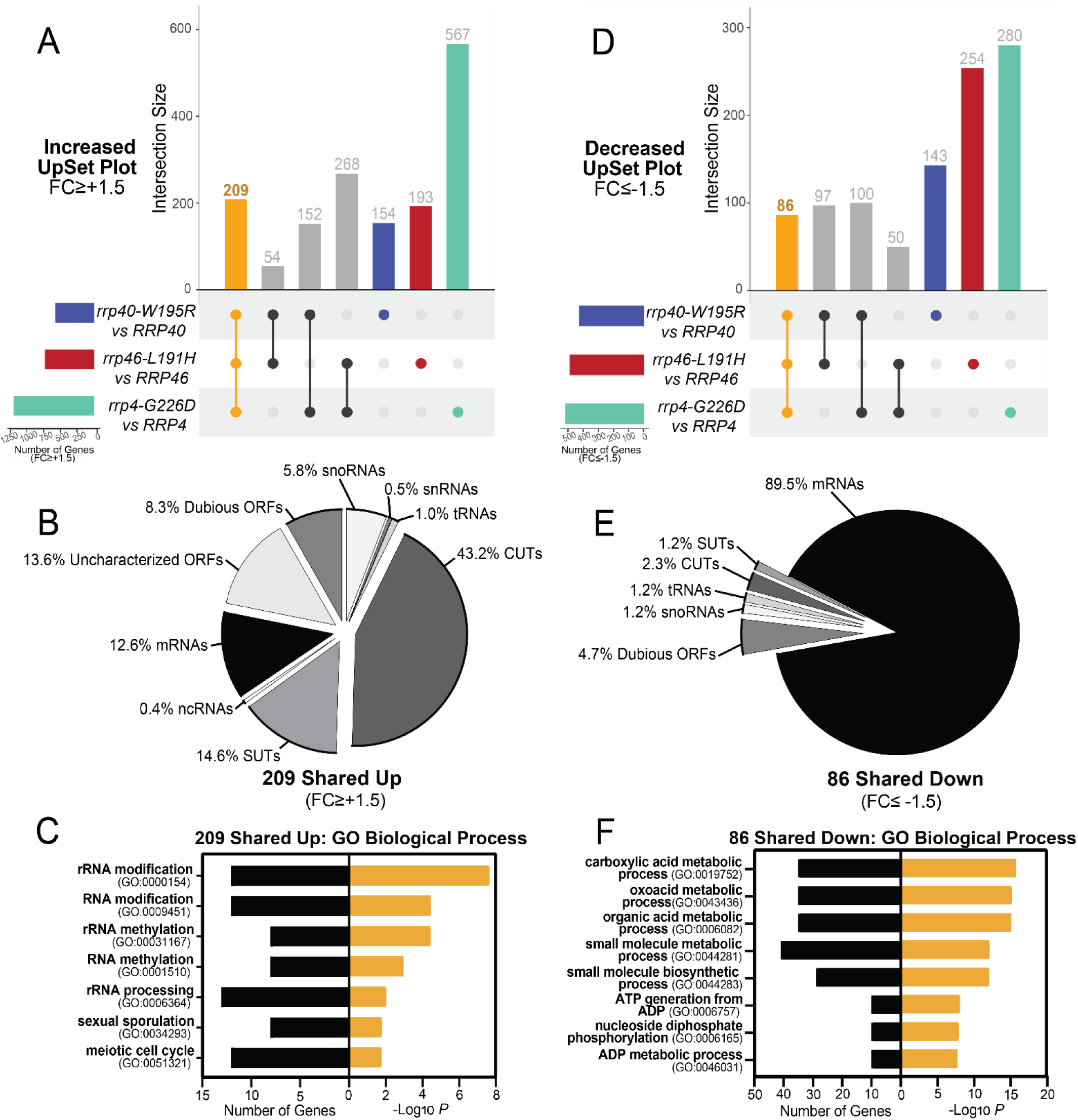
UpSet Plots of differentially expressed transcripts in *rrp4-G226D, rrp40-W195R* and *rrp46-L191H* cells reveal shared targets involved in metabolism and rRNA processing. The UpSet plot of (**A**) significantly increased (FC≥+1.5) transcripts or (**D**) significantly decreased (FC≤-1.5) transcripts within the *rrp4-G226D, rrp40-W195R* and *rrp46-L191H* datasets. Pie chart of the RNA classes that comprise the intersection of shared (**B**) increased (Up) transcripts or (**E**) decreased (Down) transcripts (Up). Gene ontology (GO) analysis for biological process of the shared (**C**) increased transcripts or (**F**) decreased transcripts. The sets of samples are color-coded: transcripts identified with FC≤-1.5 or FC≥+1.5 from differential expression analysis of *rrp40-W195R* vs *RRP40* cells are colored blue, those of *rrp46-L191H* vs *RRP46* cells are colored red, and those of *rrp4-G226D* vs *RRP4* cells are colored teal. Black bars in C and F represent the number of transcripts that are linked to each biological process category, whereas orange bars represent the-log of the associated p*-*value for each GO term. GO analyses were performed on coding RNAs (mRNA) and non-coding RNAs (tRNAs, snoRNAs, and snRNAs).

We also detected 86 transcripts shared that are significantly decreased (**Figure 4D**) among the *rrp4-G226D, rrp40-W195R* and *rrp46-L191H* cells. Of these decreased transcripts, ∼90% are mRNAs (**Figure 4E**). Gene Ontology (GO) analysis on these shared decreased transcripts revealed metabolic and biosynthetic biological processes as the top significantly impacted categories (GO:0019752; GO:0043436; GO:0006082) (**Figure 4F**). GO analyses on the identifiable human homologs of the shared increased and decreased transcripts (**Supplementary Documentation S1)** show that while most increased transcripts are yeast-specific CUTs and SUTs, those with human homologs are enriched in synaptic vesicle priming and fusion biological processes (GO:0016082; GO:0031629; GO:0099500). Similarly, the shared decreased transcripts with human homologs are enriched in metabolic and biosynthetic processes (GO:0019752; GO:0043436; GO:0006082). This analysis suggests that the modeled RNA exosomopathy mutations result in changes in highly conserved metabolic and biosynthetic pathways.

Comparing mutant pairs, we identified shared significantly increased transcripts (**Figure S4A**). Most increased transcripts shared between *rrp40-W195R* and *rrp46-L191H* are mRNAs (**Figure S4B**). GO analysis of these transcripts revealed significant enrichment in rRNA and ncRNA processing and ribosome biogenesis (GO:0016072; GO:0034470; GO:0006364; GO:0042254) (**Figure S4C**). However, shared increased transcripts in *rrp40-W195R* and *rrp4-G226D* cells mostly consist of CUTs and SUTs, with no significant biological process enrichment (**Figure S4B**). Similarly, shared increased transcripts in *rrp4-G226D* and *rrp46-L191H* cells are mostly CUTs and SUTs (**Figure S4B**), with GO analysis revealing a significant enrichment in telomere maintenance and mitotic recombination (GO:0000722; GO:0006312) (**Figure S4D**). GO analyses of the shared increased transcripts between *rrp4-G226D* and *rrp40-W195R*, and *rrp40-W195R* and *rrp46-L191H* reveal significant enrichment of biosynthetic processes (GO:0043604; GO:1901566; GO:0034645).

We also identified shared significantly decreased transcripts among the pairs of models (**Figure S4E**). Most decreased transcripts are mRNAs (**Figure S4F**), but a notable percentage of shared transcripts in the *rrp4-G226D* and *rrp46-L191H* cells are from the tRNA genes (**Figure S4F**). GO analyses on the shared decreased transcripts between paired groups reveal enrichment in different biological processes related to translation. Specifically, a significant number of the shared significantly decreased transcripts in *rrp4-G226D* and *rrp40-W195R* cells impact cytoplasmic translation (GO:0002181) (**Figure S4G**) and those in *rrp4-G226D* and *rrp40-L191H* cells show translation elongation (GO:0006414) (**Figure S4H**). This is consistent with the large percentage of tRNAs that are decreased in *rrp4-G226D* and *rrp46-L191H* cells. We found no significant enrichment of any specific biological process for the shared significantly decreased transcripts in the *rrp40-W195R* and *rrp46-L191H* cells.

In summary, transcripts involved in ribosome biogenesis are significantly enriched across all three mutants, as expected given the major role of the RNA exosome in rRNA processing. The *rrp40-W195R* and *rrp4-G226D* cells, modeling mutations in the cap subunit genes, share different targets from *rrp46-L191H* cells, modeling a mutation in a core subunit gene. These data strongly suggest that the type of RNA exosome subunit that is altered influences the specific RNA classes affected by each mutant.

### Comparative assessment of differentially expressed transcripts specific to *rrp4-G226D, rrp40-W195R* or *rrp46-L191H* mutant suggest impacts on translation and ribosome biogenesis

In examining unique changes caused by each variant, we identified 154 transcripts increased specifically in the *rrp40-W195R* cells, 193 in the *rrp46-L191H* cells, and 567 in the *rrp4-G226D* cells (**Figure 5A**). Analysis of these specific transcript sets reveals a divergent pattern between the three mutant models (**Figure 5B**). In the *rrp40-W195R* cells, nearly 25% map to snoRNA, snRNA, or tRNA genes, another 25% map to CUTs and SUTs, another 25% map to mRNAs, and the rest map to dubious or uncharacterized open reading frames (ORFs). In contrast, *rrp46-L191H*-specific increased transcripts are predominantly mRNAs, while the *rrp4-G226D-*specific increased RNAs are mostly CUTs and SUTs. GO analysis of the *rrp40-W195R*-specific transcripts revealed significant enrichment in biological processes involved in gene expression (GO:0010467), rRNA modification (GO:00000154), and translation elongation (GO: 0006414) (**Figure 5C**). For the *rrp46-L191H* cells, GO analysis revealed significant enrichment in processes related to ncRNA processing (GO:0034470) and ribosome biogenesis (GO:0042254) (**Figure 5D**). No significant enrichment was found for the *rrp4-G226D*-specific increased transcripts, likely due to the large percentage of CUTs and SUTs.

**Figure 5.**
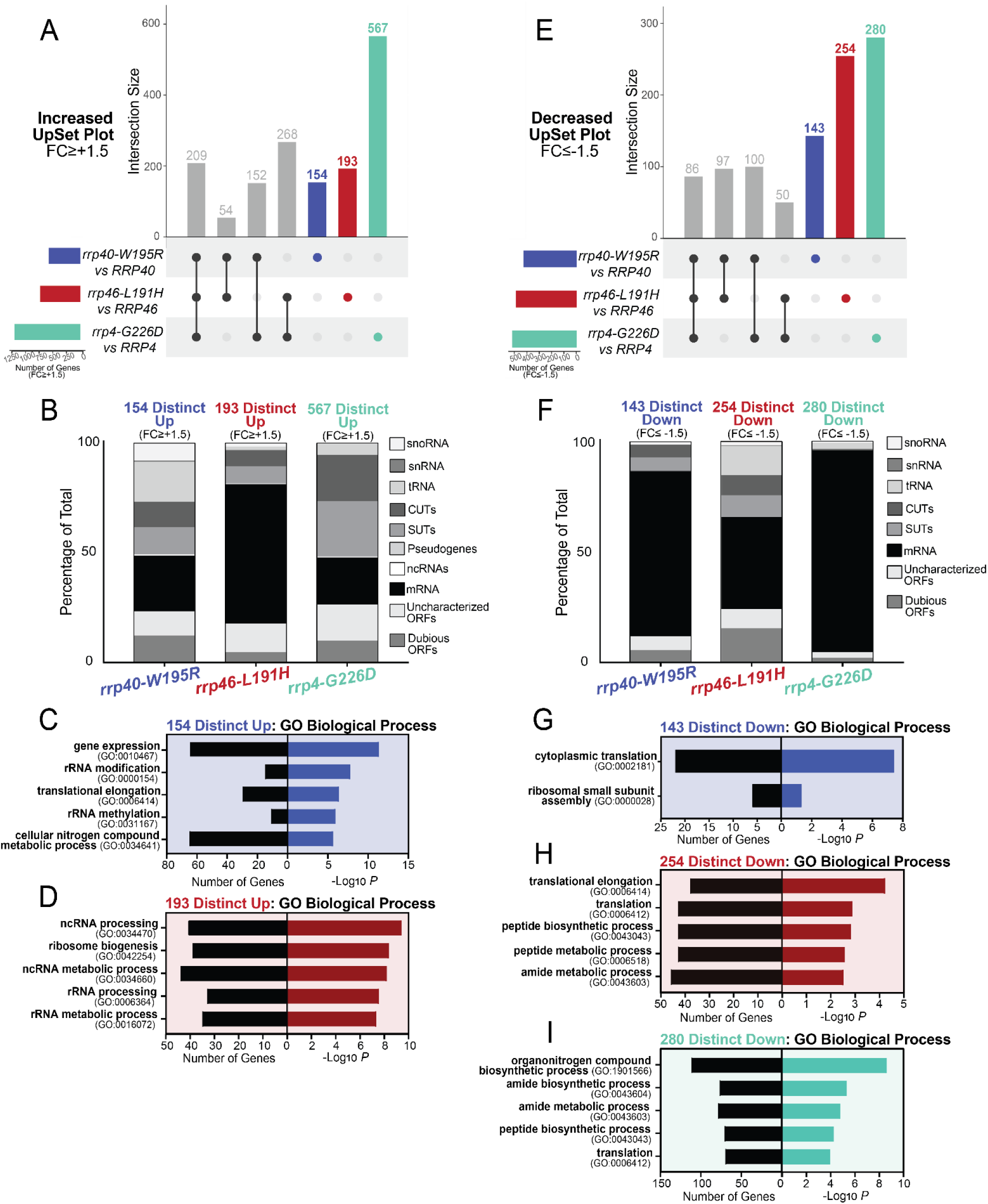
UpSet Plots of differentially expressed transcripts in *rrp4-G226D, rrp40-W195R* and *rrp46-L191H* cells reveal RNA targets that are uniquely impacted. UpSet plots were generated as in Figure 4. The intersections assessed here are transcripts significantly increased or decreased by 1.5-fold or more only in the *rrp40-W195R* dataset (blue), the *rrp46-L191H* dataset (red), or the *rrp4-G226D* dataset (teal). The UpSet plot of significantly (**A**) increased (FC≥+1.5) transcripts or (**E**) decreased (FC≤-1.5) transcripts occurring solely in the *rrp40-W195R*, *rrp46-L191H*, or *rrp4-G226D* dataset. Stacked bar percentages of the RNA types that comprise the (**B**) increased (Up) transcripts or (**F**) decreased (Down) transcripts identified only in the *rrp40-W195R, rrp46-L191H,* or *rrp4-G226D* dataset. Gene ontology (GO) analysis for biological process of the (**C**) increased transcripts or (**G**) decreased transcripts occurring only in the *rrp40-W195R* dataset; (**D**) increased transcripts or (**H**) decreased transcripts occurring only in the *rrp46-L191H* dataset, and the (**I**) decreased transcripts occurring only in the *rrp4-G226D* dataset. In panels, C-D and G-I, black bars represent the number of transcripts that are linked to each biological process category, whereas colored bars represent the-log of the associated *p-*value for each GO term. All GO analyses were performed on coding RNAs (mRNA) and non-coding RNAs (tRNAs, snoRNAs, and snRNAs).

We identified 143 transcripts significantly decreased specifically in the *rrp40-W195R* cells, 254 in the *rrp46-L191H* cells, and 280 in the *rrp4-G226D* cells (**Figure 5E**). Distinct patterns emerge from the analysis of these transcripts, with the cap mutants (*rrp4-G226D* and *rrp40-W195R*) primarily affecting mRNAs, while the *rrp46-L191H* core mutant impacts a more diverse range of transcripts, including tRNA gene transcripts, CUTs and SUTs (**Figure 5F**). GO analysis of the 143 decreased transcripts in the *rrp40-W195R* cells revealed significant enrichment in biological processes related to cytoplasmic translation (GO:0002181) and ribosomal small subunit assembly (GO:0000028) (**Figure 5G**). The 254 decreased transcripts in *rrp46-L191H* and 280 in *rrp4-G226D* are enriched in translation (GO:0006412), amide and peptide biosynthesis, and metabolic processes (**Figure 5H-I**).

Overall, the GO analyses of transcripts uniquely changed in each mutant model revealed enrichment in several similar biological processes, particularly ribosome biogenesis, translation and biosynthesis. However, the transcripts that produce the shared GO terms are distinct within each mutant cell model. These data suggest that while there may be overlapping impacts on key biological processes, the overall consequences are in part due to distinct target changes.

### RNA exosomopathy yeast mutant models exhibit defects in ribosome biogenesis, impact global translation, and alter translation fidelity

The RNA exosome is required for the processing of rRNAs (Schmid and Jensen 2008; Fernández-Pevida et al. 2015). In line with this function, GO analyses of the RNA-seq data from *rrp4-G226D, rrp40-W195R,* and *rrp46-L191H* cells revealed enrichment in terms related to translation and ribosome biogenesis (**Figures 4C** and **5C-D**). In addition, differential expression analysis revealed that some of the most significantly decreased transcripts in all three mutant models are *RPS* and *RPL* mRNAs, which encode ribosomal proteins. To broadly compare impacts on ribosomal protein genes across the *rrp4-G226D, rrp40-W195R* and *rrp46-L191H* cells, we generated heatmaps of normalized FPKM expression estimates specifically for the *RPS* and *RPL* genes (**Figure 6**), using FPKM estimates and including data from *rrp6Δ* cells collected in the same RNA-seq experiment. Consistent with previous work, there is a decrease in transcript levels for most ribosomal protein genes in *rrp6Δ* cells (Fox et al. 2015). Our analysis also included mitochondrial ribosomal protein genes (*MRPS* and *MRPL*), which in large part show a different transcriptomic pattern change, compared to *RPS* and *RPL* transcripts, in *rrp6Δ* cells. Furthermore, we detect an overall reduction of ribosomal protein gene transcripts in all three RNA exosomopathy models with the biological triplicates clustering together. However, the magnitude of the decrease is less than the decrease observed in *rrp6Δ* cells. To assess the effect of RNA exosome variants on ribosome biogenesis, we examined rRNA processing by northern blotting of total RNA (**Figure 7**). For these assays, we generated CRISPR/Cas9-edited *rrp4-G226D*, *rrp40-W195R*, and *rrp46-L191H* mutant cells to be able to compare them to each other directly and parent wild-type cells, instead of comparing each deletion with a point mutant plasmid to the same deletion with a wild-type plasmid. The temperature-sensitive growth of the *rrp4/40/46* CRISPR mutants was confirmed compared to the wild-type parental cells in a solid media growth assay, and rescue of the growth of the mutants by wild-type *RRP4*, *RRP40*, or *RRP46* plasmid was confirmed (**Figure S5**). Consistent with previously published data for *rrp4-G226D*, *rrp40-W195R, and rrp46-L191H* (Gillespie et al. 2017; Slavotinek et al. 2020; Sterrett et al. 2021), northern blots show that all RNA exosome variants cause significant accumulation of 7S pre-rRNA, a precursor of mature 5.8S rRNA that is processed by the RNA exosome (**Figure 7A, B**). The magnitude of 7S accumulation is higher for *rrp4-G226D* than for the other models (**Figure 7C**), consistent with the more severe growth defect of *rrp4-G226D* cells compared to the other mutants (**Figure 2**), resulting in a more significant reduction of mature 5.8S rRNA in these cells (**Figure 7D**). In addition, cap mutant cells (*rrp4-G226D* and *rrp40-W195R*) show a clear accumulation of 27SA2, which is not seen in the core mutant cells (*rrp46-L191H*) (**Figure 7A-B**). In contrast, *rrp46-L191H* cells show significant accumulation of 23S pre-rRNA, a precursor of 18S rRNA, accompanied by a reduction in levels of 20S rRNA (**Figure 7A**), resulting in a significant reduction of mature 18S rRNA (**Figure 7D**). The *rrp46-L191H* rRNA processing defects are similar to those of another core mutant (*rrp41-L187P*), which we recently analyzed (Fasken et al. 2024). Overall, the ribosome biogenesis defects in the cap and core mutant cells exhibit both shared and distinct characteristics. While all mutants result in accumulation of 7S pre-rRNAs, the cap mutants have a more significant impact on the biogenesis of the 60S subunit, which is also reflected in a reduced ratio of 25S/18S rRNA in these cells (**Figure 7D**).

**Figure 6.**
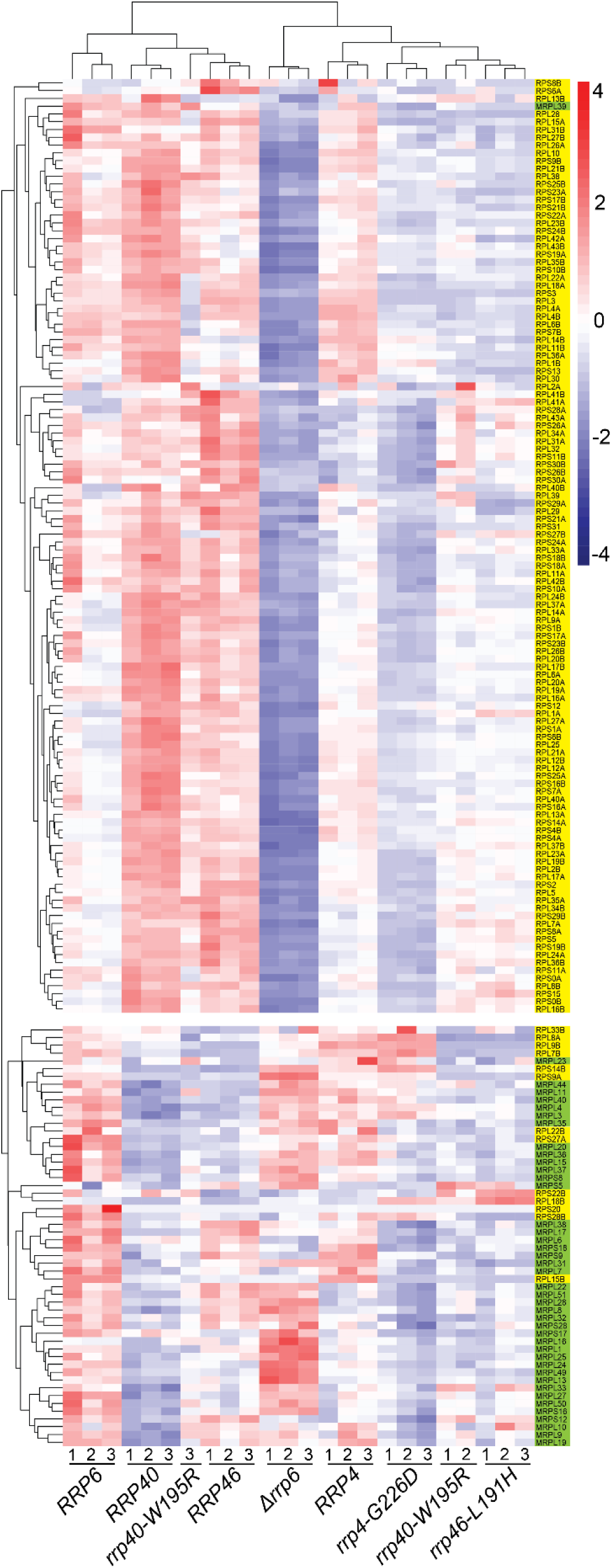
Heatmaps of RNA exosomopathy yeast mutants reveal broad changes in ribosomal protein gene transcript levels. Heatmaps were generated with FPKM estimates of all annotated ribosomal protein *RPS* and *RPL* genes. Cytoplasmic and mitochondrial subunit genes are shaded in yellow and green, respectively. Gene expression estimates are scaled for heatmap visualization, and coloring is a gradient of higher (red) to lower (blue) scaled values. The gene name for each *RPS* or *RPL* transcript is listed on the right

**Figure 7.**
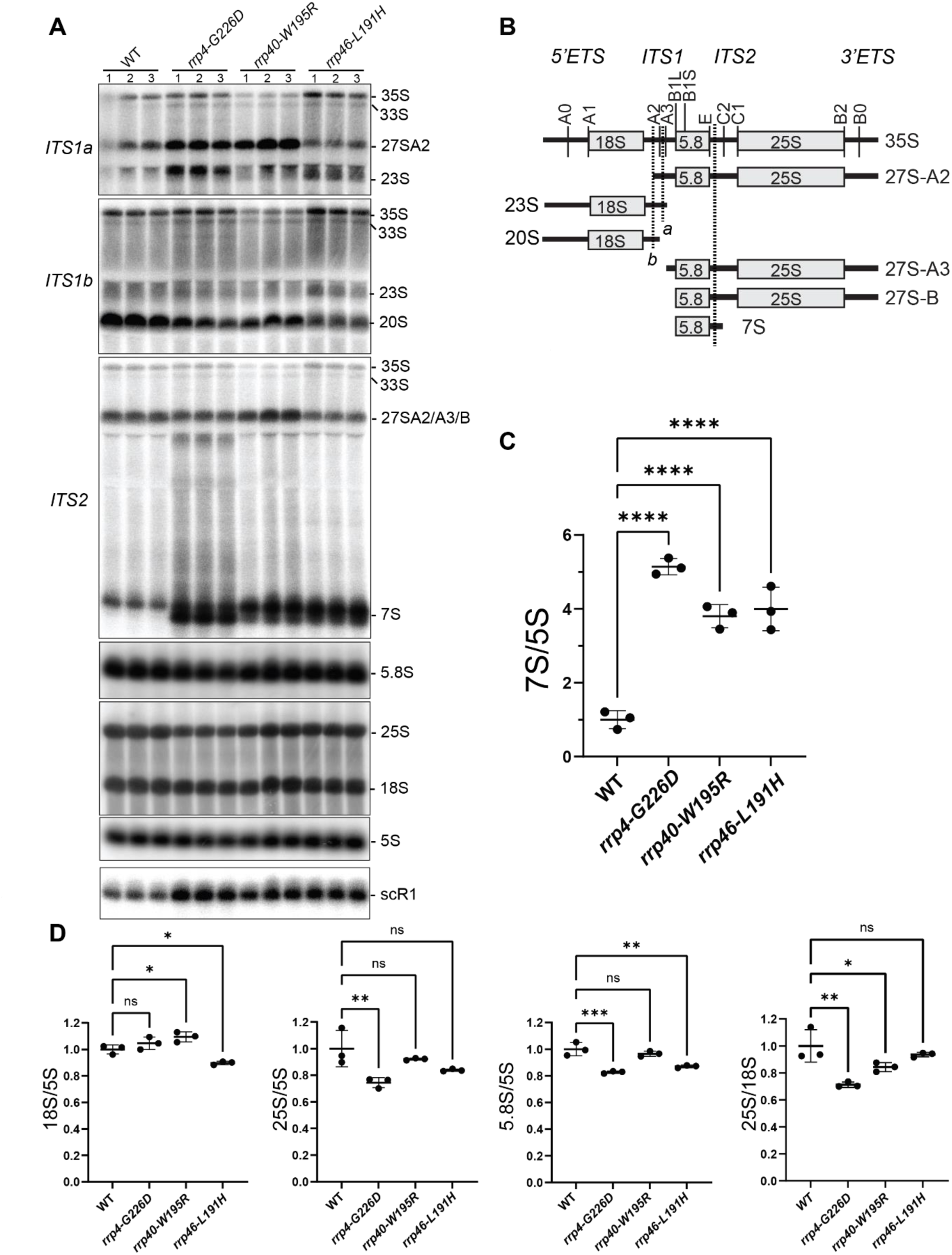
Ribosome biogenesis defects in *rrp4-G226D, rrp40-W195R,* and *rrp46-L191H* mutant cells. (**A**) Northern blot analysis of steady-state levels of precursor rRNA levels in the indicated cells. Three biological replicates were grown at 30°C and analyzed. (**B**) Schematic of the yeast precursor rRNAs and the binding site of the probes used in (**A**). Probes used to detect 18S, 25S, 5.8S, 5S and scR1 bind to an internal region within the sequence of the mature RNA. (**C**) Quantification of the ratio of 7S precursor rRNA to mature 5S rRNA for data shown in **(A**). (**D**) Quantification of 18S, 25S, and 5.8S mature rRNA levels relative to 5S rRNA, and the ratio of 25S to 18S, in the RNA exosomopathy mutant yeast models compared to wild-type control cells for three biological replicates shown in (**A**).

### Polysome profiling on RNA exosomopathy mutant yeast models reveals translation differences that suggest distinct molecular consequences in ribosome pools between cap and core mutants

To further explore how specific ribosome biogenesis defects caused by RNA exosomopathy variants affect cellular translation, we performed polysome profiling on mutant and wild-type control cells. As expected from cells with severe ribosome biogenesis defects, all three mutant cells show a significant decrease in the level of polysomes compared to wild-type control cells (**Figure 8A**). In addition, the *rrp4/40* cap subunit mutants show an accumulation of halfmer polysomes, evident as shoulders on the 80S monosome peak in the *rrp4-G226D* cells and on the monosome, disome and trisome peaks in the *rrp40-W195R* cells, not found in the profiles from the wild-type cells or *rrp46-L191H* core subunit mutant (**Figure 8A**) or another recently analyzed core subunit mutant (*rrp41-L187P*) (Fasken et al. 2024). The halfmers could form due to inefficient ribosome subunit joining during translation initiation because of defects/reduction in biogenesis or stability of 60S subunits compared to 40S. Indeed, the 25S/18S rRNA ratio is decreased in the cap mutant cells (**Figure 7D**). Additionally, the polysome profiles of the cap subunit mutants exhibit a distinct pattern of free 40S and 60S peaks when compared to those of the core subunit mutant described here and another core subunit mutant that was recently analyzed (*rrp41-L187P*) (Fasken et al. 2024), in which the free 60S peak shows a pronounced spike and free 40S subunits are reduced (**Figure 8A**). Thus, the polysome profiles in the cap *vs* core mutant yeast cells appear to be distinctly different.

**Figure 8.**
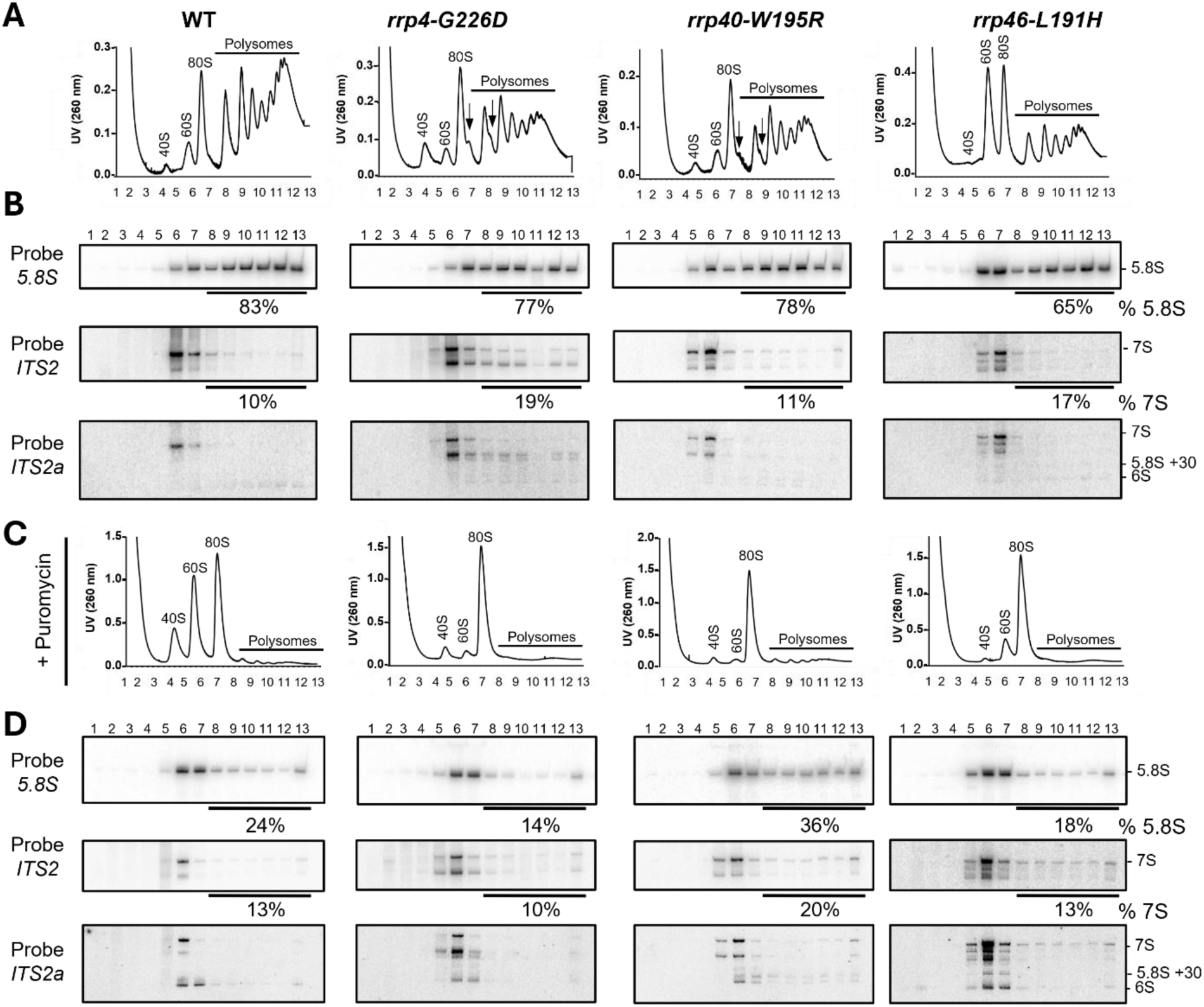
Distinct translation defects in *rrp4-G226D, rrp40-W195R,* and *rrp46-L191H* mutant cells. (**A** and **C**) Sucrose density gradients of wild-type, *rrp4-G226D*, *rrp40-W195R*, and *rrp46-L191H* cells that were grown at 30°C are shown. Clarified cell extracts were resolved on a 10-50% sucrose gradient and scanned at 260 nm. Arrows indicate halfmers. (**B** and **D**) Northern blots of gradient fractions indicating the distribution of 5.8S rRNA and 7S pre-rRNA. Samples in (**C**) and (**D**) were treated with 2.5 mM puromycin after lysis.

The Rrp4-G226D variant has decreased association with the essential RNA helicase Mtr4 (Sterrett et al. 2021). The helicase Mtr4 (budding yeast ortholog of human MTR4/MTREX) aids the RNA exosome in targeting the 5.8S rRNA precursor (7S rRNA) among other target RNA transcripts and makes direct contact with the RNA exosome cap subunit Rrp4 and cofactors Rrp6 and Mpp6 (de la Cruz et al. 1998; LaCava et al. 2005; Vaňáčová et al. 2005; Houseley and Tollervey 2008; Houseley and Tollervey 2009; Lubas et al. 2011; Stuparevic et al. 2013; Schuch et al. 2014; Rodríguez-Galán et al. 2015; Kilchert et al. 2016; Falk et al. 2017; Zinder and Lima 2017; Weick et al. 2018). Previous work has shown that a temperature-sensitive mutant of *MTR4* in budding yeast results in escape of 7S pre-rRNA-containing ribosomes into the pool of translating ribosomes (Rodríguez-Galán et al. 2015). To assess whether and how RNA exosomopathy yeast mutants affect the quality of translating ribosomes, we assayed the distribution of 7S pre-rRNA in polysome fractions of mutant cells compared to control wild-type cells (**Figure 8B**). In wild-type cells, 7S pre-rRNA peaks with the free 60S fraction, where precursor 60S subunits would co-migrate. In contrast, in *rrp4-G226D* mutant cells, a fraction of 7S pre-rRNA co-migrates with polysomes. Overall, in *rrp4-G226D* and *rrp40-W195R* cap mutants, the spread of 7S pre-rRNA in polysome fractions is broader and across all polysome fractions, compared to the 7S pre-rRNA distribution in the *rrp46-L191H* core mutant and another recently described core variant (*rrp41-L187P*) (Fasken et al. 2024). This effect is pronounced at 37°C where the cells have a more significant growth defect (**Figure S6A-B**).

To determine whether the 7S pre-rRNA containing complexes migrating in polysome fractions in *rrp4-G226D* mutant cells are simply aggregated complexes, we prepared cell lysates under polysome run-off conditions by omitting cycloheximide and adding 2.5 mM puromycin, which dissociates the 80S ribosomes into 40S and 60S subunits (Blobel and Sabatini 1971). Under these conditions, a portion of 7S pre-rRNAs and 5.8S rRNAs in *rrp4-G226D* cells was shifted to the 80S and 60S fractions in all tested strains (**Figure 8C**). Again, this effect was more pronounced at 37°C (**Figure S6C-D**). Northern blots also showed that the pattern of 7S pre-rRNA processing/degradation was distinct between the *rrp4-G226D, rrp40-W195R* and *rrp46-L191H* mutants (**Figure 8B and S6B**). Together, these data further corroborate the differential molecular consequences of the RNA exosomopathy mutants.

To test how exosomopathy mutants impact the quality of protein synthesis, we assayed translation fidelity (**Figure 9A-C**), using previously established dual-luciferase reporters (Harger and Dinman 2003; Salas-Marco and Bedwell 2005; Cheung et al. 2007). In these reporter plasmids, Renilla luciferase is constitutively expressed, whereas the production of firefly luciferase is dependent on a translation fidelity defect, such as a programmed frameshifting event, recognition of an alternative start site, or miscoding. Interestingly, the *rrp4-G226D* and *rrp40-W195R* cells show a statistically significant decrease in decoding the H245R near-cognate mutant firefly luciferase mRNA compared to wild-type controls (**Figure 9A-B**). Together, the polysome profiles combined with dual luciferase reporter assays indicate a severe defect in translation in RNA exosomopathy yeast mutant models. At least some of these defects, e.g. formation of halfmers, appear to correlate with the type of RNA exosome subunit that is changed, *i.e*., cap vs. core subunit, in the tested variants.

**Figure 9.**
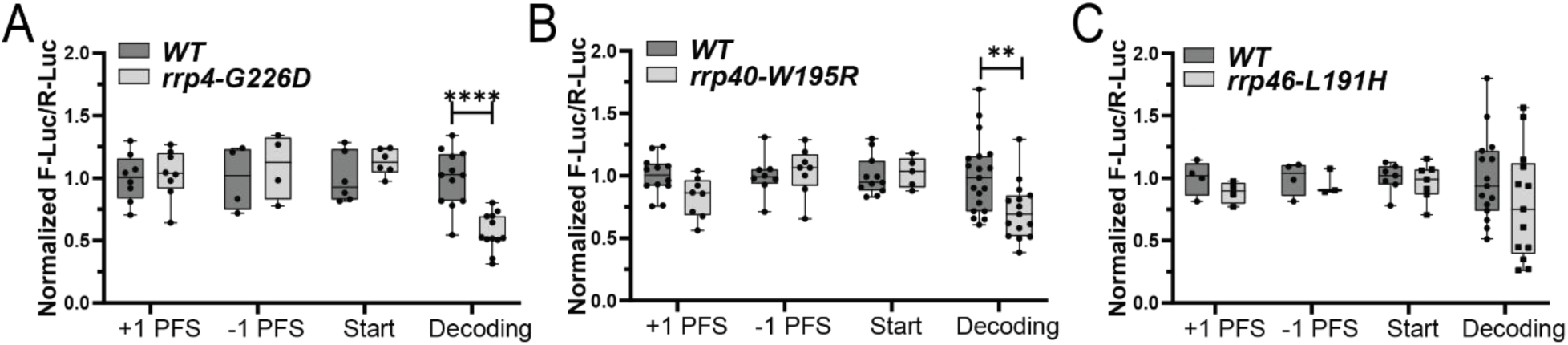
Translation fidelity defects in the RNA exosomopathy mutant yeast models compared to wild-type control cells. Dual-luciferase reporter assays were used to measure translation fidelity. The expression of firefly and Renilla luciferase was measured in *rrp4-G226D* (**A**), *rrp40-W195R* (**B**), and *rrp46-L191H* (**C**) mutant cells, compared to the corresponding wild-type control cells. The ratio of firefly luciferase to Renilla luciferase was normalized to control plasmids. Four to twelve biological replicates were analyzed.

## DISCUSSION

This work represents the first *in vivo* comparative study of a collection of RNA exosomopathy mutant models. Comparison of the cellular RNAs that show changes in steady-state levels across the modeled RNA exosomopathy mutations in yeast provides intriguing results regarding the shared and distinctly altered transcripts and the potential pathways – metabolic and biosynthetic processes, rRNA processing/modification, and ribosome biogenesis – impacted within the models analyzed. Additionally, this study explores the consequences of these RNA exosome mutants for translation, including identifying some differences in translational defects between the *rrp4-G226D* and *rrp40-W195R* cap mutants and the *rrp46-L191H* core mutant. The results of this study underscore the significance of the link between impairment of RNA exosome function and defects in translation efficiency and fidelity, providing insights into the cellular defects that may contribute to distinct pathologies reported in exosomopathies.

The majority of studies that have explored the functional consequences of mutations underlying exosomopathies have focused on a single RNA exosome subunit (Morton et al. 2018; Fasken et al. 2020). Those studies that have compared across altered subunits have employed patient cells (Burns et al. 2018), which introduces inherent variability into these comparisons. The present study takes advantage of the evolutionary conservation of the RNA exosome complex to create a series of exosomopathy models that can be compared to one another to define similarities and differences in molecular phenotypes. These studies in budding yeast will not define cell-type specific RNA targets but can provide insight into the different classes of RNAs impacted.

We acknowledge that different RNA classes (*e.g.,* tRNAs and snoRNAs compared to mRNAs) require different sequencing approaches to be comprehensively analyzed. We also note that some of the transcript differences detected will be indirect and reflect reduced growth rates and stress responses or different degrees of penetrance of the specific alleles, transcripts targeted for degradation or processing intermediates. Thus, whether the molecular consequences observed in RNA exosomopathy models are more systemic or are directly caused by changes in the integrity of the RNA exosome complex, remains to be fully established. Addressing these questions will require an in-depth biochemical analysis of each RNA exosome subunit variant to determine the impact of these changes on the RNA exosome complex and interactome. However, some predictions can be made based on the structures of the RNA exosome, considering the distinct consequences of each pathogenic RNA exosome subunit variant modeled in this study. The EXOSC2-G198D cap subunit variant is predicted to severely impact the structural organization of the cap (Di Donato et al. 2016; Sterrett et al. 2021). In contrast, the EXOSC3 cap subunit variants are predicted to impact interactions with interacting core subunits within the complex. The EXOSC3-D132 residue lies in a loop between strands in the S1 domain, and the EXOSC3-D132A variant would therefore likely be impaired in folding and interactions with neighboring subunits EXOSC5 and EXOSC9 (Fasken et al. 2017; Fasken et al. 2020). Similarly, the EXOSC3-W238 residue is predicted to position other EXOSC3 residues to interact with neighboring EXOSC9 residues. Thus, the EXOSC3-W238R variant could weaken EXOSC3•EXOSC9 interactions (Fasken et al. 2017; Fasken et al. 2020).

The EXOSC5 core subunit variants have been predicted to be unstable and impair complex interactions (Slavotinek et al. 2020). The EXOSC5-T114 residue interacts with A62 in the N-terminal region of the subunit and the EXOSC5-T114I variant would therefore likely be disrupted in this intra-subunit interaction (Slavotinek et al. 2020). The EXOSC5-M148 residue is at the interface with EXOSC3 and the EXOSC5-M148T variant would therefore likely cause impaired interactions between EXOSC5 and EXOSC3. The EXOSC5-L206 residue is buried in a hydrophobic pocket of the subunit. Therefore, the EXOSC5-L206H variant would likely cause destabilizing effects on the integrity of the subunit (Slavotinek et al. 2020). Lastly, the EXOSC9-L14 residue is located in the first alpha helix of EXOSC9 and EXOSC9-L14P variant would therefore likely cause disrupted interactions within the subunit (Liu et al. 2016). Thus, the pathogenic amino acid substitutions are predicted to have varied biochemical structural consequences. As such, each RNA exosomopathy protein variant may have differential impacts on the overall structure and function of the RNA exosome complex that require further investigation.

In growth assays, the *rrp4-G226D* mutant, modeling SHRF-linked *EXOSC2-G198D,* the *rrp40-W195R* mutant, modeling PCH-linked *EXOSC3-W238R,* and the *rrp46-L191H* mutant, modeling cerebellar atrophy-linked *EXOSC5-L206H* show the most impaired growth in budding yeast. These results are intriguing as the associated RNA exosomopathy disease pathologies linked to each mutation modeled in *S. cerevisiae* are diverse in the tissues impacted. Neither the *EXOSC3-W238R* mutation nor the *EXOSC2-G198D* mutation has been found in the homozygous state, suggesting these mutations may be lethal or highly deleterious in humans (Wan et al. 2012; Biancheri et al. 2013; Rudnik-Schoneborn et al. 2013; Eggens et al. 2014; Halevy et al. 2014; Di Donato et al. 2016). In contrast, the *EXOSC5-L206H* mutation, which has been found in the homozygous state, is associated with early infant mortality (Slavotinek et al. 2020). While the exosomopathy yeast mutant models all show defects, the modeled *rrp4-G226D, rrp40-W195R* and *rrp46-L191H* mutants can support cell viability when expressed as the sole copy of the essential *RRP* gene, allowing us to analyze the functional consequences of these pathogenic amino acid changes in the budding yeast system.

Differential expression analysis reveals a significant decrease in transcripts that encode various Heat shock protein (HSP) family members in the *rrp4-G226D* cells. The considerable reduction in *HSP* mRNAs could suggest that the *rrp4-G226D* cells have a compromised response to heat stress, thus explaining the significant growth defect observed at 37°C. Loss of Rrp6 also causes a decrease in levels of *HSP* transcripts (Wang et al. 2020), however in a manner independent of the interaction between the exonuclease and the RNA exosome complex. Whether loss of Rrp6 due to altered interaction with Rrp4 also contributes to a decrease in *HSP* mRNAs in the *rrp4-G226D* cells is not yet clear. The compromised response to stress observed in *rrp4-G226D* cells may also explain the increase in specific transcripts (e.g., *DDR2* and *PIR3* mRNAs), as expression of both these genes is activated in response to a variety of stress conditions (Gasch et al. 2001; Ribeiro et al. 2022). Furthermore, previous work has also linked RNA exosome activity to cellular integrity upon heat stress (Novačić et al. 2021). The changes in RNA exosome activity observed within all three *rrp* mutants may contribute to the slow growth observed upon heat stress. Additionally, the decrease in several biosynthetic transcripts observed in all three *rrp* mutants could further contribute to slow growth phenotypes. These changes to conserved metabolic and biosynthetic pathways observed in the model *rrp* mutants could also potentially contribute to disease pathologies in individuals with RNA exosomopathies. Among the increased transcripts detected in differential expression analysis of the three *rrp* mutant models, some had identifiable human homologs. GO analysis of the evolutionarily conserved transcripts revealed significant enrichment in synaptic vesicle priming and fusion biological processes. The link between these modeled RNA exosomopathy mutations and biological processes involved in synaptic vesicle fusion and trafficking may provide context for the numerous neurological defects common in individuals with mutations in *EXOSC* genes.

All three *rrp* mutant models analyzed in detail also show a significant decrease in a number of mRNAs. Many of these changes in mRNA levels could result from systemic effects, reflecting changes in cell signaling that can occur when the cell is under stress due to a dysfunctional RNA exosome. GO analysis of shared dysregulated transcripts reveals pathways related to ribosome biogenesis and translation are significantly enriched, pointing to potential changes in ribosome levels and/or function. In line with this observation, northern blots confirm a decrease in mature rRNA levels in *rrp4-G226D*, *rrp40-W195R,* and *rrp46-L191H* cells (Gillespie et al. 2017; Slavotinek et al. 2020). Depletion of individual RNA exosome subunits was previously shown to impact pre-rRNA cleavage, even at steps not directly involving the action of the RNA exosome, resulting in rRNA processing defects and reduction of mature rRNAs (Allmang et al. 2000). Indeed, besides involvement in 60S ribosome subunit maturation (Fernández-Pevida et al. 2015), the nuclear RNA exosome stably associates with late 90S pre-ribosomes during their transition to pre-40S and the RNA exosome is required for remodeling of 90S and maturation of pre-40S subunits (Lau et al. 2021). Our data corroborate the significant effect of these pathogenic RNA exosome mutants on ribosome biogenesis and further show that the effects of variants in specific subunits of the RNA exosome on translation goes beyond impacting solely the defined function of the RNA exosome in pre-rRNA processing. Some of these effects could be caused by accumulation or reduction of levels of transcripts that are important for ribosome assembly, including the snoRNAs and the specific mRNA transcripts coding for ribosomal proteins or ribosome assembly factors. Other effects could be direct or indirect due to reduced or incomplete processing of the precursor rRNAs. Impaired processing of rRNA could result in assembly of dysfunctional ribosomes that are either quality controlled and discarded, causing reduced ribosome numbers, or escape into the translating pool of ribosomes.

Maturation of the 3′-end of the 5.8S rRNA from the 7S pre-rRNAs involves a series of cleavage events performed by different nucleases (Fernández-Pevida et al. 2015). In budding yeast, the trimming of 7S to 6S pre-rRNA happens in the nucleus whereas the final processing of 6S pre-rRNA to 5.8S rRNA is cytoplasmic (Thomson and Tollervey 2010). However, immature 60S ribosome subunits have previously been reported to escape the nucleus and engage in translation in mutant yeast (Briggs et al. 1998; Rodríguez-Galán et al. 2015). In *rrp6Δ* cells, pre-60S ribosomes containing nuclear 5.8S rRNA and an additional 30 nucleotides of the ITS2 region can escape to the cytoplasm and enter the polysome pool (Briggs et al. 1998). Similarly, expression of a variant of the RNA exosome cofactor Mtr4 helicase was reported to result in the accumulation of nuclear pre-60S ribosomes containing 7S pre-rRNA in the cytoplasm (Rodríguez-Galán et al. 2015). These premature ribosomes were shown to engage in translation, suggesting 7S pre-rRNA processing defects do not prevent the export of 7S-containing pre-60S ribosomes (Rodríguez-Galán et al. 2015). In the context of 40S maturation, the trimming of the 3’-end of 18S rRNA from its precursor 20S pre-rRNA also occurs in the cytoplasm (Fatica et al. 2003; Fatica et al. 2004; Lamanna and Karbstein 2009; Pertschy et al. 2009), and this process is strictly quality controlled to prevent the entry of premature 20S-containing 40S ribosomal subunits into the translating pool of ribosomes (Parker et al. 2019). However, cytoplasmic pre-40S ribosomes can escape into the translating pool in several mutant yeast mutants that fail the surveillance pathways during 40S assembly (Soudet et al. 2010; Strunk et al. 2012; García-Gómez et al. 2014; Parker et al. 2019; Parker and Karbstein 2023).

Our data indicate that the cap vs core RNA exosome subunit variants cause distinct translational defects, with cap variant Rrp4-G226D and Rrp40-W195R showing formation of halfmers and indicating a potential problem in 60S maturation or subunit joining, while the core Rrp46-L191H and Rrp41-L187P (Fasken et al. 2024) variants also indicating 40S biogenesis defect features. Our results suggest that a fraction of 7S pre-rRNA ribosomes in *rrp4-G226D* cells enter the translation pool, albeit not as efficiently as the 5.8S-containing ribosomes. Future mechanistic studies are needed to dissect how a shortage of ribosomes and the presence of 7S-containing ribosomes in the translation pool impact the cellular proteome in these RNA exosomopathy models. Overall, these studies demonstrate that variants in both the cap and core RNA exosome subunits could significantly change the translating ribosome pool and cellular proteome. However, the cap and core variants are likely to impact translation in distinct ways, possibly due to the different rRNA intermediates, contributing to unique molecular consequences and pathologies in each mutant model.

Our data also show that RNA exosomopathy variants result in translation fidelity defects. These defects could arise from the lower ribosome levels (Mills and Green 2017) or changes in transcripts related to ribosome biogenesis or translation, *e.g.,* mRNAs encoding ribosome biogenesis and translation factors, tRNAs and/or snoRNAs. The decoding defect may be indicative of decreased rates of translation elongation in exosomopathy mutant cells, which likely provides more time for discrimination between aminoacyl tRNAs (Plant et al. 2007). Changes in tRNA abundance could also contribute to the decoding defect. Although we detected changes in transcripts that map to tRNAs, our sequencing approach was not ideally suited to measure tRNA abundance in an unbiased manner. Future studies are therefore required to reveal how decoding is altered in RNA exosomopathy mutant cells. Understanding whether RNA exosomopathy variants lead to changes in tRNA abundance is especially intriguing, as some RNA exosomopathies exhibit clinical features similar to those seen in patients with pontocerebellar hypoplasias caused by mutations of the tRNA splicing endonuclease (TSEN) subunit genes (Hayne et al. 2023).

Of the two cap subunit variants modeled, the *rrp4-G226D* variant causes a severe growth defect at 37°C, whereas the *rrp40-W195R* variant causes a slight growth defect at 37°C. Thus, a global shortage of RNA exosome complexes cannot simply explain the severity of the molecular impact from each mutant or the overall growth fitness. Therefore, even though the simplest model to explain the severity of RNA exosomopathy mutants may be an RNA exosome concentration decrease model that would affect target transcript levels, the molecular pathology of RNA exosomopathy mutants appears far more complex and likely impacted by transcriptional, post-transcriptional and translational mechanisms that require further investigation.

In summary, the data presented here provide novel insights into the molecular mechanisms of defects caused by disease-causing mutations in the RNA exosome genes modeled in yeast and reveal that different RNA exosomopathy mutations result in both unique and shared molecular changes across the variants. Our data suggest that while RNA exosomopathies result from a conglomeration of many direct and systemic molecular changes at the transcript level, translational defects could also contribute to the distinct cellular outcomes. Thus, even though distinct with respect to the molecular changes, RNA exosomopathies share phenotypic and mechanistic similarities among themselves and with ribosomopathies. Future work is required to reveal why, despite affecting molecular mechanisms that are seemingly universal to all cells, specific populations of neuronal cells are most affected by RNA exosomopathies. Likely, changes in the transcript levels, ribosome numbers, quality, and fidelity of translation in RNA exosomopathies alter the translation of specific mRNAs involved in neuronal differentiation and/or function to cause tissue-specific molecular defects. These studies in yeast models provide initial mechanistic insights and suggest models to explore in relevant cell types in the future.

## MATERIALS AND METHODS

*Saccharomyces cerevisiae strains and plasmids. S. cerevisiae* strains and plasmids used in this study are listed in **Table S2**. Oligonucleotides used for plasmid construction are listed in **Table S3**. The *rrp4*Δ (yAV1103), *rrp40*Δ (yAV1107), and *rrp46*Δ (yAV1105) strains used in this study were previously described (Schaeffer et al. 2009; Slavotinek et al. 2020). The *rrp45*Δ (yAV1410) strain used in this study was obtained by transforming a *RRP45 URA3* plasmid into the heterozygous diploid *RRP45/rrp45Δ* strain from the knockout collection and sporulating the transformants to obtain a haploid *rrp45Δ* strain. The *rrp6*Δ strain (ACY1641) was obtained from Horizon Discovery. The wild-type *RRP4* (pAC3656), *RRP40* (pAC3652), and *RRP46* (pAC3482) plasmids were constructed as previously described and each contain the open reading frame (ORF) flanked by endogenous regulatory sequences (promoter, terminator and 5’/3’ UTR) cloned into the pRS315 vector (ATTC #77144) (Sikorski and Hieter 1989), which harbors the *LEU2* marker (Fasken et al. 2017; Slavotinek et al. 2020; Sterrett et al. 2021). The wild-type *RRP45* (pAV975) plasmid contains the ORF flanked by endogenous regulatory sequences cloned into the pRS415 *LEU2* vector. The *rrp4-G226D* (pAC3659), *rrp40-S87A* (pAC3654), *rrp40-W195R* (pAC3655), *rrp46-Q86I* (pAC3483), *rrp46-L127T* (pAC3534), and *rrp46-L191H* (pAC3484) mutant plasmids encoding the different RNA exosomopathy amino acid variants were generated by site-directed mutagenesis of the respective wild-type plasmids (pAC3656, pAC3652, pAC3482) using the QuikChange II Site-Directed Mutagenesis Kit (Agilent) and oligonucleotides containing the desired missense mutations as previously described (Fasken et al. 2017; Slavotinek et al. 2020; Sterrett et al. 2021). The *rrp45-I15P* (pAC3480) mutant plasmid was generated by site-directed mutagenesis of pAV975 using oligonucleotides AC8001 and AC8002 in this study. The wild-type *RRP6* (pAC3752) plasmid was constructed by subcloning ApaI/SacI-digested *RRP6* ORF with endogenous regulatory sequences from pAC2301 (*RRP6* in pRS313; Fasken et al. PLOS Genetics (2015)) into pRS315 cut with ApaI/SacI. The *TEF1p-Cas9-CYC1t-SNR52p* (pAC3846) pCas9 plasmid was constructed by PCR amplification of *TEF1p* with oligonucleotides AC8410 and AC8411 and *Cas9-CYC1t* with AC6802 and AC6803 using p414-*TEF1p-Cas9-CYC1t* plasmid template (Addgene #43802, (DiCarlo et al. 2013)) and *SNR52p* with AC6804 and AC6805 using p426-SNR52p-gRNA.CAN1.Y-SUP4t plasmid template (Addgene #43803, (DiCarlo et al. 2013)), followed by sequential cloning of SacI/SpeI-digested *TEF1p*, SpeI/KpnI-digested *Cas9-CYC1t*, and AgeI/KpnI-digested *SNR52p* into pRS316 plasmid digested with corresponding restriction enzymes. The *TEF1p-Cas9-CYC1t-SNR52p-RRP4_668.gRNA-SUP4t* (pAC3863), *TEF1p-Cas9-CYC1t-SNR52p-RRP40_583.gRNA-SUP4t* (pAC3861), and *TEF1p-Cas9-CYC1t-SNR52p-RRP46_589.gRNA-SUP4t* (pAC4342) pCas9 plasmids containing the gRNAs for targeting *RRP4*, *RRP40*, and *RRP46*, respectively, were constructed by PCR amplification of *RRP4_668.gRNA* with oligonucleotides AC8407 and AC6809, *RRP40_583.gRNA* with AC8402 and AC6809, and *RRP46_589.gRNA* with AC9888 and AC6809 using p426-SNR52p-gRNA.CAN1.Y-SUP4t plasmid template (Addgene #43803) and cloning of SphI/KpnI-digested gRNA products into pAC3846 digested with SphI/KpnI. All plasmids were fully sequenced.

*Generation of S. cerevisiae mutant strains* using *CRISPR-Cas9 genome editing*. The *rrp4-G226D* (ACY3110), *rrp40-W195R* (ACY3117), and *rrp46-L191H* (ACY3137) mutant strains were generated using CRISPR/Cas9 editing with a single pCas9-gRNA expression plasmid and double-stranded homology-directed repair (HDR) oligonucleotides in a wild-type BY4741 strain essentially as described before (DiCarlo et al. 2013). The single pCas9-gRNA plasmid on a pRS316 (*URA3, CEN6*) backbone is derived from p414-*TEF1p-Cas9-CYC1t* plasmid (Addgene #43802) and p426-SNR52p-gRNA.CAN1.Y-SUP4t plasmid (Addgene #43803). The constitutive expression of Cas9 is driven by the *TEF1* promoter and constitutive expression of the gRNA is driven by the *SNR52* promoter. Specifically, 500 ng of pAC3846 (pCas9 without gRNA), pAC3863 (pCas9 + *RRP4* gRNA) +/-1 nmol of double-stranded *rrp4-G226D* HDR oligonucleotide (AC8408/8409), pAC3861 (pCas9 + *RRP40* gRNA) +/-1 nmol double-stranded *rrp40-W195R* HDR oligonucleotide (AC8404/8405), or pAC4342 (pCas9 + *RRP46* gRNA) +/-1 nmol double-stranded *rrp46-L191H* HDR oligonucleotide (AC9889/9890) and 50 µg salmon sperm DNA was transformed into wild-type BY4741 cells by standard Lithium Acetate (LiOAc) transformation protocol (Da et al. 2000). HDR oligonucleotides are listed in **Table S3**. Cells were plated on SD-Ura media plates and incubated at 30°C for 2 days. Large colonies on plates with cells transformed pCas9-gRNA and HDR oligonucleotides were restreaked to new SD-Ura media plates and screened for the presence of *rrp4-G226D*, *rrp40-W195R*, and *rrp46-L191H* mutations via Sanger sequencing of genomic *RRP4*, *RRP40*, and *RRP46* PCR products, respectively.

*S. cerevisiae transformations and growth assays.* All yeast transformations were performed using the standard Lithium Acetate (LiOAc) protocol (Da et al. 2000). Standard plasmid shuffle assays were performed to assess the *in vivo* function of the *rrp* variants as previously described (Fasken et al. 2017; Slavotinek et al. 2020; Sterrett et al. 2021). The *rrpΔ* cells (*rrp4Δ* (yAV1103), *rrp40Δ* (yAV1107), *rrp45Δ* (yAV1410), *rrp46Δ* (yAV1105)) transformed with the wild-type control *LEU2* plasmid (*RRP4* (pAC3656), *RRP40* (pAC3652), *RRP45* (pAV975), *RRP46* (pAC3482)) or the mutant variant plasmid (*rrp4-G226D* (pAC3659), *rrp40-S87A* (pAC3654), *rrp40-W195R* (pAC3655), *rrp45-I15P* (pAC3480), *rrp46-Q86I* (pAC3483), *rrp46-L127T* (pAC3534), *rrp46-L191H* (pAC3484)) were streaked on 5-FOA SD-Leu media plates and incubated at 30°C for 2-3 days. Single colonies from the 5-FOA SD-Leu media plates were selected in quadruplicate and streaked onto selective SD-Leu media plates. The *rrp6Δ* cells (ACY1641) were transformed with empty *LEU2* vector (pRS315) or wild-type *LEU2* plasmid (*RRP6* (pAC3752)) and selected on SD-Leu media plates. The *rrpΔ* cells containing only the wild-type *RRP* or mutant *rrp LEU2* plasmid were used for the RNA-seq analysis and reporter assays. The CRISPR/Cas9-edited *rrp4-G226D* (ACY3110), *rrp40-W195R* (ACY3117), and *rrp46-L191H* (ACY3137) mutant cells were assessed for temperature-sensitive (ts) growth by a solid media growth assay and rescue of the ts growth of the mutant cells by wild-type *RRP4*, *RRP40*, and *RRP46* plasmids was confirmed. The *rrp4-G226D*, *rrp40-W195R*, and *rrp46-L191H* CRISPR mutant cells were used for the northern blots and the polysome profiling assay. Growth assays were performed on solid media and in liquid culture. The wild-type control and mutant model cells were grown to saturation at 30°C before the concentrations were adjusted to an A_600_ ∼0.5, and samples were serially diluted in 10-fold steps and spotted onto SD-Leu media plates. Plates were grown at 30°C and 37°C for 2-3 days. For growth in liquid culture, saturated overnight cultures grown at 30°C were diluted to an A_600_ ∼0.01 in SD-Leu in a 24-well plate, and growth at 37°C was monitored and recorded at OD_600_ in a BioTek® SynergyMx microplate reader with Gen5™ v2.04 software over 24 hr. For each sample analyzed in growth assays at least 3 independent biological replicates were used. In addition, for the liquid culture assays, technical triplicates for each biological sample were grown. Doubling times were calculated using GraphPad Prism version 9.3.1.

*Sample collection for RNA-seq analysis.* RNA-seq was performed on three independent biological replicates of *rrp*Δ cells containing the *RRP* wild-type control plasmids or the *rrp* variants as the sole copy of the RNA exosome gene. The *rrp6*Δ cells (ACY1641) contained either an *RRP6* wild-type control plasmid (pAC3752) or an empty vector. Biological replicates of all samples were first screened by solid media growth assays prior to growth and collection for the RNA-seq experiment. For sample collection, cells were grown in SD-Leu media overnight at 30°C to saturation, diluted to A_600_ ∼0.2 in SD-Leu media and shifted to 37°C for 5 hours. Cells were washed, pelleted and flash frozen and cell pellets were sent to Zymo Research for total RNA preparation and RNA-seq analysis.

*RNA-seq Library Preparation.* RNA-seq library preparation was performed by Zymo Research. Total RNA-seq libraries were constructed from 300 ng of total RNA. To remove rRNA, a method previously described (Bogdanova et al. 2011) was followed with some modifications. Libraries were prepared using the Zymo-Seq RiboFree Total RNA Library Prep Kit (Cat # R3000). RNA-seq libraries were sequenced on an Illumina NovaSeq to a sequencing depth of at least 20 million read pairs (150 bp paired-end sequencing) per sample.

*Sequence Data Alignments and Differential Expression Analysis.* NovaSeq paired-end 150-bp reads from Total RNA-seq data files were first adaptor trimmed and then analyzed using the STAR program (version 2.6.1d) for alignment of short reads to *S. cerevisiae* reference genome. Transcript and gene expression estimates were measured using StringTie v2.1.7 (Pertea et al. 2015). The expression estimates fragments per Kilobase of transcript per Million mapped (FPKM) were used with the Pheatmap R package v1.0.12 to generate heatmaps (Kolde 2012). The raw reads per gene were counted using featureCounts v1.22.2 (Liao et al. 2013) to the *S. cerevisiae* S288C genome assembly R64-1-1 (Engel et al. 2014), annotated with CUTs and SUTs (Xu et al. 2009). Low feature counts (<10 reads total) were removed. Differential gene expression analysis on raw read counts was performed using the DESeq2 R package v1.38.1 (Love et al. 2014) to identify genes significantly changed (*p*-value<0.05, ≥1.5-fold change) in *rrp* mutant variant samples relative to *RRP* wild-type control samples. Shrinkage of effect size was performed on differential expression data for visualizations using the *apeglm* method (Zhu et al. 2019). Using DESeq2, principal component analysis (PCA) was performed, and MA plots were generated on raw read counts. Volcano plots of differential gene expression data were produced using EnhancedVolcano R package v1.16.0 (Blighe et al. 2019). UpSet plots were generated using UpSetR R package v1.4.0 (Conway et al. 2017), with transcripts identified through differential expression analysis in the mutant cells as significantly decreased by 1.5-fold or more (FC ≤-1.5) and significantly increased by 1.5-fold or more (FC ≥ +1.5). UpSet intersections are shown in graphs as a matrix, with rows corresponding to the sets of samples (i.e., transcripts identified FC ≤-1.5 or FC ≥ +1.5 within the three mutant models) and columns corresponding to the intersection between these sets. Pie charts and stacked bars of RNA class percentages in significantly altered genes were generated using GraphPad Prism version 9.3.1. Transcripts were sorted by class using the annotations available through the *Saccharomyces cerevisiae Genome Database* (SGD) (Cherry et al. 2012). Gene Ontology (GO) analysis on significantly altered genes for Biological Process category was performed using the YeastMine webserver. GO analysis on human homolog genes was performed using HumanMine. All GO analyses were performed by Holm-Bonferroni test correction.

*Northern Blot Analysis of rRNAs.* For analysis of ribosomal RNAs, yeast cells were grown to mid-log phase and RNA was extracted using the hot phenol method. Northern blotting was carried out essentially as previously described (Khoshnevis et al. 2019), using the probes listed in **Table S3**.

*Dual-Luciferase Reporter Assays to Monitor Translation Fidelity Defects.* For assaying translation fidelity defects, cells expressing wild-type or variant RNA exosome subunits and a dual-luciferase reporter plasmid were grown to mid-log phase in Ura^-^, Leu^-^ synthetic glucose liquid media. Cells were pelleted, washed, and stored at −80°C before analysis. Luciferase activities were measured using the Dual-Luciferase Reporter Assay kit (Promega). Thawed frozen cell pellets were resuspended in 1 X Passive Lysis Buffer and incubated for 10 min. LARII was mixed with lysate in clear bottom 96-well Microplates (Costar), and Firefly luciferase activity was measured. Stop and Glo solution was added, and Renilla luciferase activity was measured. Measurements were performed using a Synergy Microplate reader (BioTek). For each biological replicate (single transformant), the Firefly luciferase signal was normalized to the Renilla luciferase signal. For each strain, Firefly/Renilla ratio was normalized to the average Firefly/Renilla ratio of replicates containing a control plasmid.

*Sucrose density gradient analysis.* To analyze polysomes by gradients, cells were grown to mid-log phase in YP-dextrose media at 30_°_C and 37_°_C and harvested after adding 0.1 mg/ml cycloheximide or no cycloheximide. Cells were washed and lysed in ice-cold gradient buffer (20 mM HEPES, pH 7.4, 5 mM MgCl_2_, 100 mM NaCl, and 2 mM DTT) supplemented with complete protease Inhibitor cocktail (Roche) and 0.1 mg/ml cycloheximide or no cycloheximide. Cells were broken by cryogenic grinding and cell lysate was cleared by centrifugation at 10,000 g for 10 min. The absorbance of cleared lysate was measured at UV_260_ and an equal amount of lysate was applied to 10–50% sucrose gradients in the gradient buffer for all samples. To samples lacking cycloheximide, 2.5 mM puromycin was added to the cleared cell lysate, incubated on ice for 15 min, and then placed at 30_°_C or 37_°_C before loading onto the gradient. Gradients were centrifuged for 2 h at 40,000 RPM in a SW41Ti rotor and fractionated using a BioComp fraction collector.

## DATA DEPOSITION

The raw RNA-seq data from this study have been submitted to the NCBI Gene Expression Omnibus (GEO) under accession GSE246957.

## Supporting information

Supplementary Documentation S1

## ACKNOWLEDGEMENTS

We thank the Corbett and Ghalei laboratory members for critical discussions and input. We thank Dr. Benjamin Barwick for his contributions in assisting with the analysis of the CoMMpass dataset. This work was supported by NIH awards R35GM138123 to H.G., R35GM141710 to AvH, and R01GM130147 to AvH and AHC, as well as funds from a Synergy II Nexus Award provided by the Woodruff Health Sciences Center (WHSC), Emory School of Medicine, the Office of the Provost, and Emory College of Arts and Sciences (ECAS) to H.G. and A.H.C. M.C.S. was supported by an F31 fellowship (GM134649) from the National Institutes of Health. L.A.C. was supported by NIH T32GM149422 and F31GM157874.

## Notes

### Competing Interest Statement

The authors have declared no competing interest.

### Summary of Updates

This version of the manuscript has been revised to reflect new data collected for the northern blotting shown in Figure 7. We also include additional controls for the polysome analyses shown in Figure 8.

## REFERENCES

Allmang C, Kufel J, Chanfreau G, Mitchell P, Petfalski E, Tollervey D. 1999a. Functions of the exosome in rRNA, snoRNA and snRNA synthesis. Embo Journal 18: 5399–5410.

Allmang C, Mitchell P, Petfalski E, Tollervey D. 2000. Degradation of ribosomal RNA precursors by the exosome. Nucleic Acids Res 28: 1684–1691.

Allmang C, Petfalski E, Podtelejnikov A, Mann M, Tollervey D, Mitchell P. 1999b. The yeast exosome and human PM-Scl are related complexes of 3’--> 5’ exonucleases. Genes Dev 13: 2148–2158.

Belair C, Sim S, Wolin SL. 2018. Noncoding RNA Surveillance: The Ends Justify the Means. Chem Rev 118: 4422–4447.

Biancheri R, Cassandrini D, Pinto F, Trovato R, Di Rocco M, Mirabelli-Badenier M, Pedemonte M, Panicucci C, Trucks H, Sander T et al. 2013. EXOSC3 mutations in isolated cerebellar hypoplasia and spinal anterior horn involvement. Journal of neurology 260: 1866–1870.

Bizzari S, Hamzeh AR, Mohamed M, Al-Ali MT, Bastaki F. 2020. Expanded PCH1D phenotype linked to EXOSC9 mutation. Eur J Med Genet 63: 103622.

Blighe K, Rana S, Lewis M. 2019. EnhancedVolcano: Publication-ready volcano plots with enhanced colouring and labeling. R package version 1: 10.18129.

Blobel G, Sabatini D. 1971. Dissociation of mammalian polyribosomes into subunits by puromycin. Proc Natl Acad Sci U S A 68: 390–394.

Boczonadi V, Mueller JS, Pyle A, Munkley J, Dor T, Quartararo J, Ferrero I, Karcagi V, Giunta M, Polvikoski T et al. 2014. EXOSC8 mutations alter mRNA metabolism and cause hypomyelination with spinal muscular atrophy and cerebellar hypoplasia. Nature Communications 5.

Bogdanova EA, Barsova EV, Shagina IA, Scheglov A, Anisimova V, Vagner LL, Lukyanov SA, Shagin DA. 2011. Normalization of Full-Length-Enriched cDNA. In cDNA Libraries: Methods and Applications, (ed. C Lu, J Browse, JG Wallis), pp. 85-98. Humana Press, Totowa, NJ.

Boivin V, Deschamps-Francoeur G, Couture S, Nottingham RM, Bouchard-Bourelle P, Lambowitz AM, Scott MS, Abou-Elela S. 2018. Simultaneous sequencing of coding and noncoding RNA reveals a human transcriptome dominated by a small number of highly expressed noncoding genes. Rna 24: 950–965.

Bonneau F, Basquin J, Ebert J, Lorentzen E, Conti E. 2009. The yeast exosome functions as a macromolecular cage to channel RNA substrates for degradation. Cell 139: 547–559.

Briggs MW, Burkard KT, Butler JS. 1998. Rrp6p, the yeast homologue of the human PM-Scl 100-kDa autoantigen, is essential for efficient 5.8 S rRNA 3’ end formation. J Biol Chem 273: 13255–13263.

Brouze M, Szpila M, Czerwińska AM, Antczak W, Mroczek S, Kuliński T, Hojka-Osińska AM, Cysewski D, Gewartowska O, Adamska D et al. 2025. DIS3L, cytoplasmic exosome catalytic subunit, is essential for development but not cell viability in mice. Rna.

Burns D, Donkervoort D, Bharucha-Goebel D, Giunta M, Munro B, Scavina M, Foley R, Müller J, Bönnemann C, Horvath R. 2017. A recessive mutation in EXOSC9 causes abnormal RNA metabolism resulting in a novel form of cerebellar hypoplasia/atrophy with early motor neuronopathy. Neuromuscul Disord 27: S38.

Burns DT, Donkervoort S, Müller JS, Knierim E, Bharucha-Goebel D, Faqeih EA, Bell SK, AlFaifi AY, Monies D, Millan F et al. 2018. Variants in EXOSC9 Disrupt the RNA Exosome and Result in Cerebellar Atrophy with Spinal Motor Neuronopathy. Am J Hum Genet 102: 858–873.

Chekanova JA, Gregory BD, Reverdatto SV, Chen H, Kumar R, Hooker T, Yazaki J, Li P, Skiba N, Peng Q et al. 2007. Genome-wide high-resolution mapping of exosome substrates reveals hidden features in the Arabidopsis transcriptome. Cell 131: 1340–1353.

Cherry JM, Hong EL, Amundsen C, Balakrishnan R, Binkley G, Chan ET, Christie KR, Costanzo MC, Dwight SS, Engel SR et al. 2012. Saccharomyces Genome Database: the genomics resource of budding yeast. Nucleic Acids Res 40: D700–705.

Cheung YN, Maag D, Mitchell SF, Fekete CA, Algire MA, Takacs JE, Shirokikh N, Pestova T, Lorsch JR, Hinnebusch AG. 2007. Dissociation of eIF1 from the 40S ribosomal subunit is a key step in start codon selection in vivo. Genes Dev 21: 1217–1230.

Conway JR, Lex A, Gehlenborg N. 2017. UpSetR: an R package for the visualization of intersecting sets and their properties. Bioinformatics 33: 2938–2940.

Corbett AH. 2018. Post-transcriptional regulation of gene expression and human disease. Curr Opin Cell Biol 52: 96–104.

Cramer P. 2019. Organization and regulation of gene transcription. Nature 573: 45–54.

Da B, Dawson D, Stearns T. 2000. Methods In Yeast Genetics: A Cold Spring Harbor Laboratory Course Manual.

Damseh NS, Obeidat AN, Ahammed KS, Al-Ashhab M, Awad MA, van Hoof A. 2023. Pontocerebellar hypoplasia associated with p.Arg183Trp homozygous variant in EXOSC1 gene: A case report. Am J Med Genet A 191: 1923-1928.

Davis CA, Ares M, Jr. 2006. Accumulation of unstable promoter-associated transcripts upon loss of the nuclear exosome subunit Rrp6p in Saccharomyces cerevisiae. Proc Natl Acad Sci U S A 103: 3262–3267.

de la Cruz J, Kressler D, Tollervey D, Linder P. 1998. Dob1p (Mtr4p) is a putative ATP-dependent RNA helicase required for the 3’ end formation of 5.8S rRNA in Saccharomyces cerevisiae. Embo j 17: 1128–1140.

Di Donato N, Neuhann T, Kahlert A-K, Klink B, Hackmann K, Neuhann I, Novotna B, Schallner J, Krause C, Glass IA et al. 2016. Mutations in EXOSC2 are associated with a novel syndrome characterised by retinitis pigmentosa, progressive hearing loss, premature ageing, short stature, mild intellectual disability and distinctive gestalt. Journal of Medical Genetics 53: 419–425.

DiCarlo JE, Norville JE, Mali P, Rios X, Aach J, Church GM. 2013. Genome engineering in Saccharomyces cerevisiae using CRISPR-Cas systems. Nucleic Acids Res 41: 4336–4343.

Eggens VRC, Barth PG, Niermeijer J-MF, Berg JN, Darin N, Dixit A, Fluss J, Foulds N, Fowler D, Hortobágyi T et al. 2014. EXOSC3 mutations in pontocerebellar hypoplasia type 1: novel mutations and genotype-phenotype correlations. Orphanet Journal of Rare Diseases 9: 23.

Engel SR, Dietrich FS, Fisk DG, Binkley G, Balakrishnan R, Costanzo MC, Dwight SS, Hitz BC, Karra K, Nash RS et al. 2014. The reference genome sequence of Saccharomyces cerevisiae: then and now. G3 (Bethesda) 4: 389-398.

Falk S, Bonneau F, Ebert J, Kogel A, Conti E. 2017. Mpp6 Incorporation in the Nuclear Exosome Contributes to RNA Channeling through the Mtr4 Helicase. Cell Reports 20: 2279–2286.

Fasken MB, Leung SW, Cureton LA, Al-Awadi M, Al-Kindy A, van Hoof A, Khoshnevis S, Ghalei H, Al-Maawali A, Corbett AH. 2024. A biallelic variant of the RNA exosome gene, EXOSC4, associated with neurodevelopmental defects impairs RNA exosome function and translation. J Biol Chem 300: 107571.

Fasken MB, Losh JS, Leung SW, Brutus S, Avin B, Vaught JC, Potter-Birriel J, Craig T, Conn GL, Mills-Lujan K et al. 2017. Insight into the RNA Exosome Complex Through Modeling Pontocerebellar Hypoplasia Type 1b Disease Mutations in Yeast. Genetics 205: 221–237.

Fasken MB, Morton DJ, Kuiper EG, Jones SK, Leung SW, Corbett AH. 2020. The RNA Exosome and Human Disease. Methods in molecular biology (Clifton, NJ) 2062: 3–33.

Fatica A, Oeffinger M, Dlakić M, Tollervey D. 2003. Nob1p is required for cleavage of the 3’ end of 18S rRNA. Mol Cell Biol 23: 1798–1807.

Fatica A, Tollervey D, Dlakić M. 2004. PIN domain of Nob1p is required for D-site cleavage in 20S pre-rRNA. Rna 10: 1698–1701.

Fernández-Pevida A, Kressler D, de la Cruz J. 2015. Processing of preribosomal RNA in Saccharomyces cerevisiae. Wiley Interdiscip Rev RNA 6: 191–209.

Fox MJ, Gao H, Smith-Kinnaman WR, Liu Y, Mosley AL. 2015. The Exosome Component Rrp6 Is Required for RNA Polymerase II Termination at Specific Targets of the Nrd1-Nab3 Pathway. PLoS genetics 11: e1004999.

García-Gómez JJ, Fernández-Pevida A, Lebaron S, Rosado IV, Tollervey D, Kressler D, de la Cruz J. 2014. Final pre-40S maturation depends on the functional integrity of the 60S subunit ribosomal protein L3. PLoS genetics 10: e1004205.

Gasch AP, Huang M, Metzner S, Botstein D, Elledge SJ, Brown PO. 2001. Genomic expression responses to DNA-damaging agents and the regulatory role of the yeast ATR homolog Mec1p. Mol Biol Cell 12: 2987–3003.

Gerlach P, Schuller JM, Bonneau F, Basquin J, Reichelt P, Falk S, Conti E. 2018. Distinct and evolutionary conserved structural features of the human nuclear exosome complex. eLife 7.

Gillespie A, Gabunilas J, Jen JC, Chanfreau GF. 2017. Mutations of EXOSC3/Rrp40p associated with neurological diseases impact ribosomal RNA processing functions of the exosome in S. cerevisiae. Rna 23: 466–472.

Gudipati RK, Xu Z, Lebreton A, Seraphin B, Steinmetz LM, Jacquier A, Libri D. 2012. Extensive degradation of RNA precursors by the exosome in wild-type cells. Mol Cell 48: 409–421.

Halevy A, Lerer I, Cohen R, Kornreich L, Shuper A, Gamliel M, Zimerman BE, Korabi I, Meiner V, Straussberg R et al. 2014. Novel EXOSC3 mutation causes complicated hereditary spastic paraplegia. Journal of neurology 261: 2165–2169.

Harger JW, Dinman JD. 2003. An in vivo dual-luciferase assay system for studying translational recoding in the yeast Saccharomyces cerevisiae. Rna 9: 1019–1024.

Hayne CK, Sekulovski S, Hurtig JE, Stanley RE, Trowitzsch S, van Hoof A. 2023. New insights into RNA processing by the eukaryotic tRNA splicing endonuclease. J Biol Chem 299: 105138.

Hou D, Ruiz M, Andrulis ED. 2012. The ribonuclease Dis3 is an essential regulator of the developmental transcriptome. Bmc Genomics 13.

Houseley J, Tollervey D. 2008. The nuclear RNA surveillance machinery: the link between ncRNAs and genome structure in budding yeast? Biochim Biophys Acta 1779: 239–246.

Houseley J, Tollervey D. 2009. The many pathways of RNA degradation. Cell 136: 763–776.

Khoshnevis S, Dreggors RE, Hoffmann TFR, Ghalei H. 2019. A conserved Bcd1 interaction essential for box C/D snoRNP biogenesis. J Biol Chem 294: 18360–18371.

Kilchert C, Wittmann S, Vasiljeva L. 2016. The regulation and functions of the nuclear RNA exosome complex. Nat Rev Mol Cell Biol 17: 227–239.

Kiss DL, Andrulis ED. 2010. Genome-wide analysis reveals distinct substrate specificities of Rrp6, Dis3, and core exosome subunits. Rna 16: 781–791.

Kolde R. 2012. Pheatmap: pretty heatmaps. R package version 1: 726.

Kowalinski E, Kögel A, Ebert J, Reichelt P, Stegmann E, Habermann B, Conti E. 2016. Structure of a Cytoplasmic 11-Subunit RNA Exosome Complex. Mol Cell 63: 125–134.

LaCava J, Houseley J, Saveanu C, Petfalski E, Thompson E, Jacquier A, Tollervey D. 2005. RNA degradation by the exosome is promoted by a nuclear polyade nylation complex. Cell 121: 713–724.

Lamanna AC, Karbstein K. 2009. Nob1 binds the single-stranded cleavage site D at the 3’-end of 18S rRNA with its PIN domain. Proc Natl Acad Sci U S A 106: 14259–14264.

Lau B, Cheng J, Flemming D, La Venuta G, Berninghausen O, Beckmann R, Hurt E. 2021. Structure of the Maturing 90S Pre-ribosome in Association with the RNA Exosome. Mol Cell 81: 293–303.e294.

Liao Y, Smyth GK, Shi W. 2013. featureCounts: an efficient general purpose program for assigning sequence reads to genomic features. Bioinformatics 30: 923–930.

Lim SJ, Boyle PJ, Chinen M, Dale RK, Lei EP. 2013. Genome-wide localization of exosome components to active promoters and chromatin insulators in Drosophila. Nucleic Acids Res 41: 2963–2980.

Liu JJ, Niu CY, Wu Y, Tan D, Wang Y, Ye MD, Liu Y, Zhao W, Zhou K, Liu QS et al. 2016. CryoEM structure of yeast cytoplasmic exosome complex. Cell Res 26: 822–837.

Liu Q, Greimann JC, Lima CD. 2006. Reconstitution, activities, and structure of the eukaryotic RNA exosome. Cell 127: 1223–1237.

Lorentzen E, Dziembowski A, Lindner D, Seraphin B, Conti E. 2007. RNA channelling by the archaeal exosome. Embo Reports 8: 470–476.

Love MI, Huber W, Anders S. 2014. Moderated estimation of fold change and dispersion for RNA-seq data with DESeq2. Genome Biology 15: 550.

Lubas M, Christensen MS, Kristiansen MS, Domanski M, Falkenby LG, Lykke-Andersen S, Andersen JS, Dziembowski A, Jensen TH. 2011. Interaction profiling identifies the human nuclear exosome targeting complex. Mol Cell 43: 624–637.

Makino DL, Baumgaertner M, Conti E. 2013. Crystal structure of an RNA-bound 11-subunit eukaryotic exosome complex. Nature 495: 70–75.

Makino DL, Schuch B, Stegmann E, Baumgärtner M, Basquin C, Conti E. 2015. RNA degradation paths in a 12-subunit nuclear exosome complex. Nature 524: 54–58.

Marin-Vicente C, Domingo-Prim J, Eberle AB, Visa N. 2015. RRP6/EXOSC10 is required for the repair of DNA double-strand breaks by homologous recombination. Journal of Cell Science 128: 1097–1107.

Mills EW, Green R. 2017. Ribosomopathies: There’s strength in numbers. Science 358.

Mitchell P, Petfalski E, Shevchenko A, Mann M, Tollervey D. 1997. The exosome: A conserved eukaryotic RNA processing complex containing multiple 3’->5’ exoribonucleases. Cell 91: 457–466.

Mitchell P, Petfalski E, Tollervey D. 1996. The 3’ end of yeast 5.8S rRNA is generated by an exonuclease processing mechanism. Genes Dev 10: 502–513.

Molleston JM, Sabin LR, Moy RH, Menghani SV, Rausch K, Gordesky-Gold B, Hopkins KC, Zhou R, Jensen TH, Wilusz JE et al. 2016. A conserved virus-induced cytoplasmic TRAMP-like complex recruits the exosome to target viral RNA for degradation. Genes & Development 30: 1658–1670.

Morton DJ, Jalloh B, Kim L, Kremsky I, Nair RJ, Nguyen KB, Rounds JC, Sterrett MC, Brown B, Le T et al. 2020. A Drosophila model of Pontocerebellar Hypoplasia reveals a critical role for the RNA exosome in neurons. PLoS genetics 16: e1008901.

Morton DJ, Kuiper EG, Jones SK, Leung SW, Corbett AH, Fasken MB. 2018. The RNA exosome and RNA exosome-linked disease. Rna 24: 127–142.

Novačić A, Beauvais V, Oskomić M, Štrbac L, Dantec AL, Rahmouni AR, Stuparević I. 2021. Yeast RNA exosome activity is necessary for maintaining cell wall stability through proper protein glycosylation. Mol Biol Cell 32: 363–375.

Parker MD, Collins JC, Korona B, Ghalei H, Karbstein K. 2019. A kinase-dependent checkpoint prevents escape of immature ribosomes into the translating pool. PLoS Biol 17: e3000329.

Parker MD, Karbstein K. 2023. Quality control ensures fidelity in ribosome assembly and cellular health. J Cell Biol 222.

Parker R. 2012. RNA Degradation in Saccharomyces cerevisae. Genetics 191: 671–702.

Pefanis E, Wang J, Rothschild G, Lim J, Chao J, Rabadan R, Economides AN, Basu U. 2014. Noncoding RNA transcription targets AID to divergently transcribed loci in B cells. Nature 514: 389–393.

Pertea M, Pertea GM, Antonescu CM, Chang T-C, Mendell JT, Salzberg SL. 2015. StringTie enables improved reconstruction of a transcriptome from RNA-seq reads. Nature Biotechnology 33: 290–295.

Pertschy B, Schneider C, Gnädig M, Schäfer T, Tollervey D, Hurt E. 2009. RNA helicase Prp43 and its co-factor Pfa1 promote 20 to 18 S rRNA processing catalyzed by the endonuclease Nob1. J Biol Chem 284: 35079–35091.

Plant EP, Nguyen P, Russ JR, Pittman YR, Nguyen T, Quesinberry JT, Kinzy TG, Dinman JD. 2007. Differentiating between near-and non-cognate codons in Saccharomyces cerevisiae. PLoS One 2: e517.

Preker P, Nielsen J, Kammler S, Lykke-Andersen S, Christensen MS, Mapendano CK, Schierup MH, Jensen TH. 2008. RNA exosome depletion reveals transcription upstream of active human promoters. Science 322: 1851–1854.

Ribeiro RA, Bourbon-Melo N, Sá-Correia I. 2022. The cell wall and the response and tolerance to stresses of biotechnological relevance in yeasts. Front Microbiol 13: 953479.

Rodríguez-Galán O, García-Gómez JJ, Kressler D, de la Cruz J. 2015. Immature large ribosomal subunits containing the 7S pre-rRNA can engage in translation in Saccharomyces cerevisiae. RNA Biol 12: 838–846.

Rudnik-Schoneborn S, Senderek J, Jen JC, Houge G, Seeman P, Puchmajerova A, Graul-Neumann L, Seidel U, Korinthenberg R, Kirschner J et al. 2013. Pontocerebellar hypoplasia type 1: clinical spectrum and relevance of EXOSC3 mutations. Neurology 80: 438–446.

Sakamoto M, Iwama K, Sekiguchi F, Mashimo H, Kumada S, Ishigaki K, Okamoto N, Behnam M, Ghadami M, Koshimizu E et al. 2021. Novel EXOSC9 variants cause pontocerebellar hypoplasia type 1D with spinal motor neuronopathy and cerebellar atrophy. J Hum Genet 66: 401–407.

Salas-Marco J, Bedwell DM. 2005. Discrimination between defects in elongation fidelity and termination efficiency provides mechanistic insights into translational readthrough. J Mol Biol 348: 801–815.

Schaeffer D, Tsanova B, Barbas A, Reis FP, Dastidar EG, Sanchez-Rotunno M, Arraiano CM, van Hoof A. 2009. The exosome contains domains with specific endoribonuclease, exoribonuclease and cytoplasmic mRNA decay activities. Nat Struct Mol Biol 16: 56–62.

Schmid M, Jensen TH. 2008. The exosome: a multipurpose RNA-decay machine. Trends Biochem Sci 33: 501–510.

Schneider C, Kudla G, Wlotzka W, Tuck A, Tollervey D. 2012. Transcriptome-wide Analysis of Exosome Targets. Molecular Cell 48: 422–433.

Schneider C, Tollervey D. 2013. Threading the barrel of the RNA exosome. Trends in Biochemical Sciences 38: 485–493.

Schottmann G, Picker-Minh S, Schwarz J, Gill E, Rodenburg R, Stenzel W, M. Kaindl A, Schuelke M. 2017. Recessive mutation in EXOSC3 associates with mitochondrial dysfunction and pontocerebellar hypoplasia.

Schuch B, Feigenbutz M, Makino DL, Falk S, Basquin C, Mitchell P, Conti E. 2014. The exosome-binding factors Rrp6 and Rrp47 form a composite surface for recruiting the Mtr4 helicase. Embo Journal 33: 2829–2846.

Schuller JM, Falk S, Fromm L, Hurt E, Conti E. 2018. Structure of the nuclear exosome captured on a maturing preribosome. Science 360: 219–222.

Sikorski RS, Hieter P. 1989. A system of shuttle vectors and yeast host strains designed for efficient manipulation of DNA in Saccharomyces cerevisiae. Genetics 122: 19–27.

Slavotinek A, Misceo D, Htun S, Mathisen L, Frengen E, Foreman M, Hurtig JE, Enyenihi L, Sterrett MC, Leung SW et al. 2020. Biallelic variants in the RNA exosome gene EXOSC5 are associated with developmental delays, short stature, cerebellar hypoplasia and motor weakness. Human molecular genetics 29: 2218–2239.

Sloan KE, Schneider C, Watkins NJ. 2012. Comparison of the yeast and human nuclear exosome complexes. Biochem Soc Trans 40: 850–855.

Somashekar PH, Kaur P, Stephen J, Guleria VS, Kadavigere R, Girisha KM, Bielas S, Upadhyai P, Shukla A. 2021. Bi-allelic missense variant, p.Ser35Leu in EXOSC1 is associated with pontocerebellar hypoplasia. Clin Genet 99: 594-600.

Soudet J, Gélugne JP, Belhabich-Baumas K, Caizergues-Ferrer M, Mougin A. 2010. Immature small ribosomal subunits can engage in translation initiation in Saccharomyces cerevisiae. Embo j 29: 80–92.

Sterrett MC, Enyenihi L, Leung SW, Hess L, Strassler SE, Farchi D, Lee RS, Withers ES, Kremsky I, Baker RE et al. 2021. A budding yeast model for human disease mutations in the EXOSC2 cap subunit of the RNA exosome complex. RNA 27: 1046–1067.

Strunk BS, Novak MN, Young CL, Karbstein K. 2012. A translation-like cycle is a quality control checkpoint for maturing 40S ribosome subunits. Cell 150: 111–121.

Stuparevic I, Mosrin-Huaman C, Hervouet-Coste N, Remenaric M, Rahmouni AR. 2013. Cotranscriptional Recruitment of RNA Exosome Cofactors Rrp47p and Mpp6p and Two Distinct Trf-Air-Mtr4 Polyadenylation (TRAMP) Complexes Assists the Exonuclease Rrp6p in the Targeting and Degradation of an Aberrant Messenger Ribonucleoprotein Particle (mRNP) in Yeast. Journal of Biological Chemistry 288: 31816–31829.

Thomson E, Tollervey D. 2010. The final step in 5.8S rRNA processing is cytoplasmic in Saccharomyces cerevisiae. Mol Cell Biol 30: 976–984.

Vaňáčová Š, Wolf J, Martin G, Blank D, Dettwiler S, Friedlein A, Langen H, Keith G, Keller W. 2005. A New Yeast Poly(A) Polymerase Complex Involved in RNA Quality Control. PLOS Biology 3: e189.

Wan J, Yourshaw M, Mamsa H, Rudnik-Schoeneborn S, Menezes MP, Hong JE, Leong DW, Senderek J, Salman MS, Chitayat D et al. 2012. Mutations in the RNA exosome component gene EXOSC3 cause pontocerebellar hypoplasia and spinal motor neuron degeneration. Nature Genetics 44: 704–U134.

Wang C, Liu Y, DeMario SM, Mandric I, Gonzalez-Figueroa C, Chanfreau GF. 2020. Rrp6 Moonlights in an RNA Exosome-Independent Manner to Promote Cell Survival and Gene Expression during Stress. Cell Reports 31: 107754.

Wasmuth EV, Januszyk K, Lima CD. 2014. Structure of an Rrp6-RNA exosome complex bound to poly(A) RNA. Nature 511: 435–439.

Wasmuth EV, Zinder JC, Zattas D, Das M, Lima CD. 2017. Structure and reconstitution of yeast Mpp6-nuclear exosome complexes reveals that Mpp6 stimulates RNA decay and recruits the Mtr4 helicase. eLife 6: 24.

Webster SF, Ghalei H. 2023. Maturation of small nucleolar RNAs: from production to function. RNA Biol 20: 715–736.

Weick EM, Puno MR, Januszyk K, Zinder JC, DiMattia MA, Lima CD. 2018. Helicase-Dependent RNA Decay Illuminated by a Cryo-EM Structure of a Human Nuclear RNA Exosome-MTR4 Complex. Cell 173: 1663–1677.e1621.

Wolin SL, Maquat LE. 2019. Cellular RNA surveillance in health and disease. Science 366: 822–827.

Wyers F, Rougemaille M, Badis G, Rousselle J-C, Dufour M-E, Boulay J, Régnault B, Devaux F, Namane A, Séraphin B et al. 2005. Cryptic Pol II Transcripts Are Degraded by a Nuclear Quality Control Pathway Involving a New Poly(A) Polymerase. Cell 121: 725–737.

Xu Z, Wei W, Gagneur J, Perocchi F, Clauder-Münster S, Camblong J, Guffanti E, Stutz F, Huber W, Steinmetz LM. 2009. Bidirectional promoters generate pervasive transcription in yeast. Nature 457: 1033–1037.

Zhu A, Ibrahim JG, Love MI. 2019. Heavy-tailed prior distributions for sequence count data: removing the noise and preserving large differences. Bioinformatics 35: 2084–2092.

Zinder JC, Lima CD. 2017. Targeting RNA for processing or destruction by the eukaryotic RNA exosome and its cofactors. Genes & Development 31: 88–100.

Zinder JC, Wasmuth EV, Lima CD. 2016. Nuclear RNA Exosome at 3.1 Å Reveals Substrate Specificities, RNA Paths, and Allosteric Inhibition of Rrp44/Dis3. Mol Cell 64: 734–745.

