## Supplementary Documentation S1 for "Comparative analyses of disease-linked missense mutations in the RNA exosome modeled in budding yeast reveal distinct functional consequences in translation"

### Supporting Information

**Included Supporting Information:** Supplementary **Figures S1-6**, **Tables S1-S3**, and **Supplemental Documentation S1**.

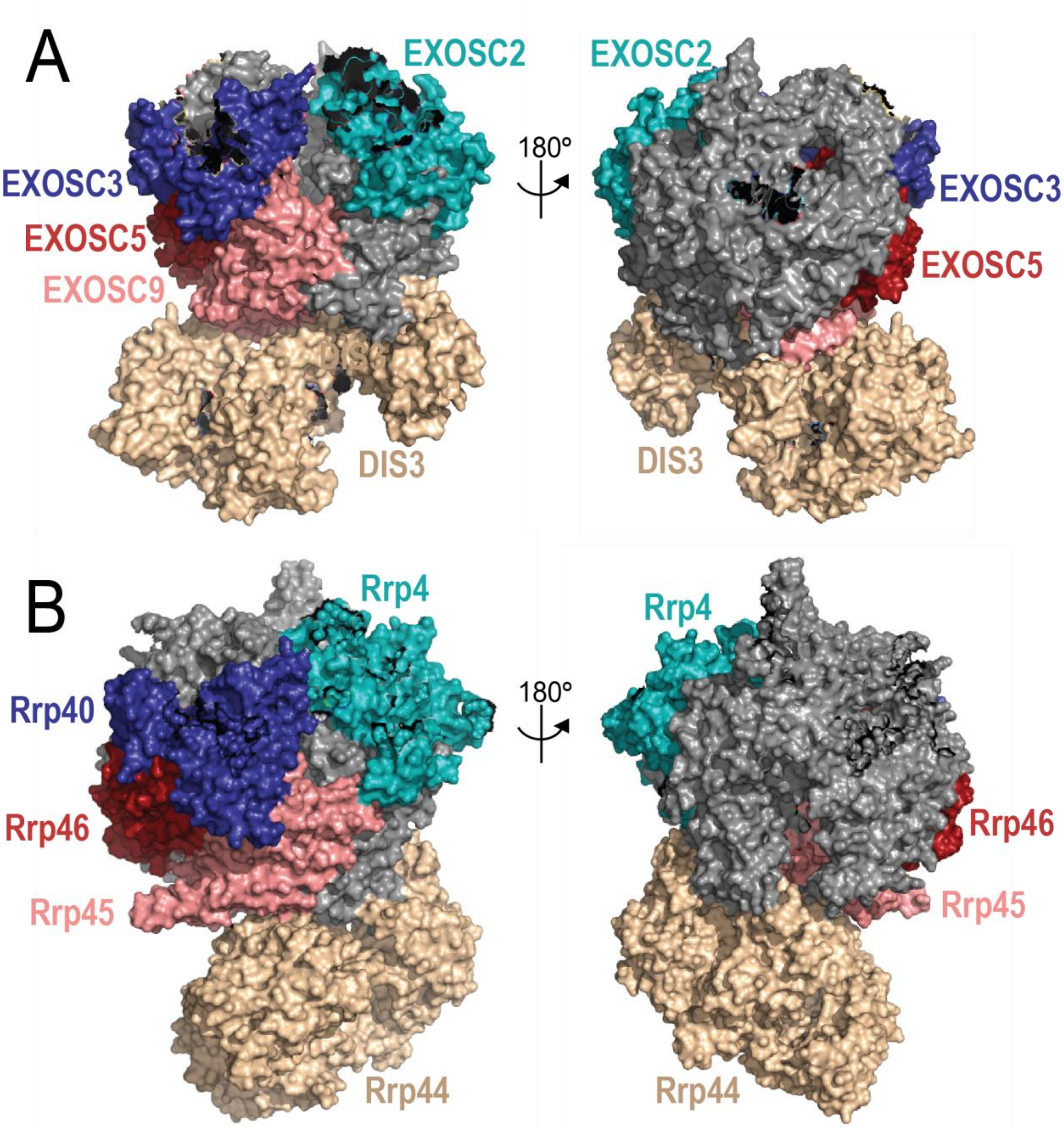


**Figure S1. The structure and organization of the RNA exosome is highly conserved across eukaryotes.** A structural model of the human RNA exosome (A) [PDB 6D6Q] and the *S. cerevisiae* RNA exosome (B) [PDB 6FSZ] are depicted with the core (Human EXOSC5/EXOSC9; *S. cerevisiae* Rrp46/Rrp45) and cap subunits (EXOSC2/EXOSC3; Rrp4/Rrp40) that are linked to RNA exosomopathy diseases labeled and color-coded.

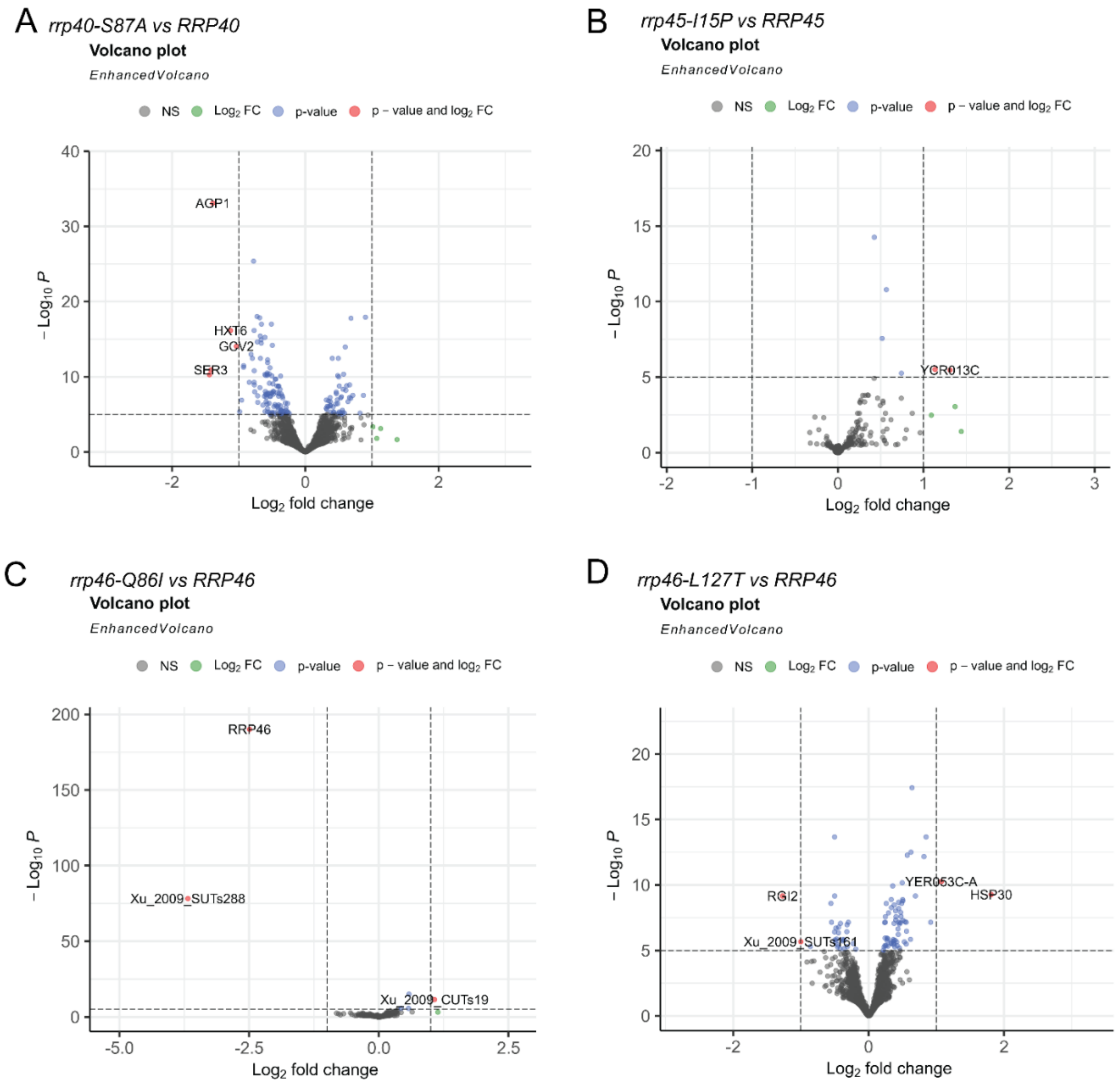


**Figure S2. Volcano plots of differentially expressed transcripts identified in *rrp40-S87A, rrp45-I15P, rrp46-Q86I* and *rrp46-L127T* samples show few significant changes.** Volcano plots of differentially expressed transcripts between each mutant sample compared to its corresponding wild-type control. Grey represents transcripts that are not significantly different between the mutant and control; green represents transcripts that are identified as increased or decreased by 1.5-fold change or more in the mutant compared to the control; blue represents transcripts that are identified as significantly different ($p<0.05$) in the mutant compared to the control; red represents transcripts that are identified as both significantly different ($p<0.05$) and increased or decreased by 1.5-fold change or more. **(A)** Volcano plot of differentially expressed transcripts in *rrp40-S87A* compared to *RRP40* wild-type control. **(B)** Volcano plot of differentially expressed transcripts in *rrp45-I15P* compared to *RRP45*. **(C)** Volcano plot of differentially expressed transcripts in *rrp46-Q86I* compared to *RRP46.* **(D)** Volcano plot of differentially expressed transcripts in *rrp46-L127T* compared to *RRP46*.

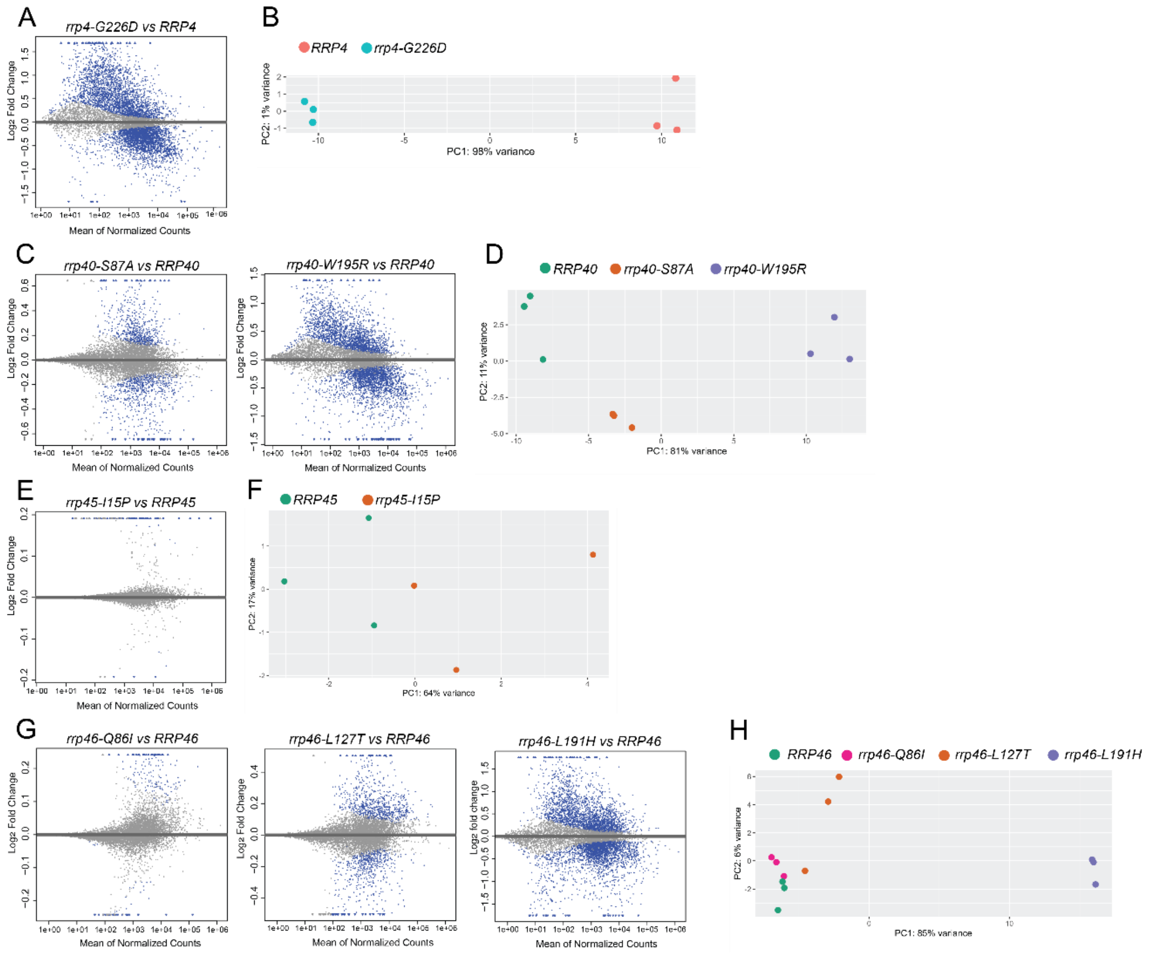


**Figure S3. MA plots for differential expression analysis results and PCA plots of sample clustering in RNA-Seq experiment.** MA plots were generated on DESeq2 results and show the log2 Fold Change (FC) values attributable to each transcript over the mean of normalized gene counts. Significant ($p<0.05$) data points are colored in blue. Principal component analysis (PCA) of RNA-seq data collected from triplicate mutant or wildtype samples is also indicated. (**A**) MA plots comparing *rrp4-G226D* to wild-type control *RRP4* samples. (**B**) PCA analysis reveals clustering of the *rrp4-G226D* samples away from the *RRP4* samples. (**C**) MA plots comparing *rrp40* mutants to wild-type control *RRP40* samples. (**D**) PCA analysis reveals clustering of the *rrp40* mutant samples away from the *RRP40* samples. The *rrp40-W195R* sample however clusters the furthest from the *RRP40* samples, encompassing 81% variance. (**E**) MA plots comparing *rrp45-I15P* to wild-type control *RRP4* samples. (**F**) PCA analysis reveals that the *rrp45-I15P* mutant samples do not independently cluster away from the *RRP45* samples, suggesting that gene expression patterns of these genotypes are not distinct. (**G**) MA plots comparing *rrp46* mutants to wild-type control *RRP40* samples. (**H**) PCA analysis reveals clustering of the *rrp46-L191H* mutant samples away from the *RRP40* samples. The *rrp46-Q86I* and *rrp46-L127T* samples do not cluster independently from the *RRP46* samples, suggesting the gene expression patterns of these *rrp46* mutants are very similar to that of the wild-type control genotype.

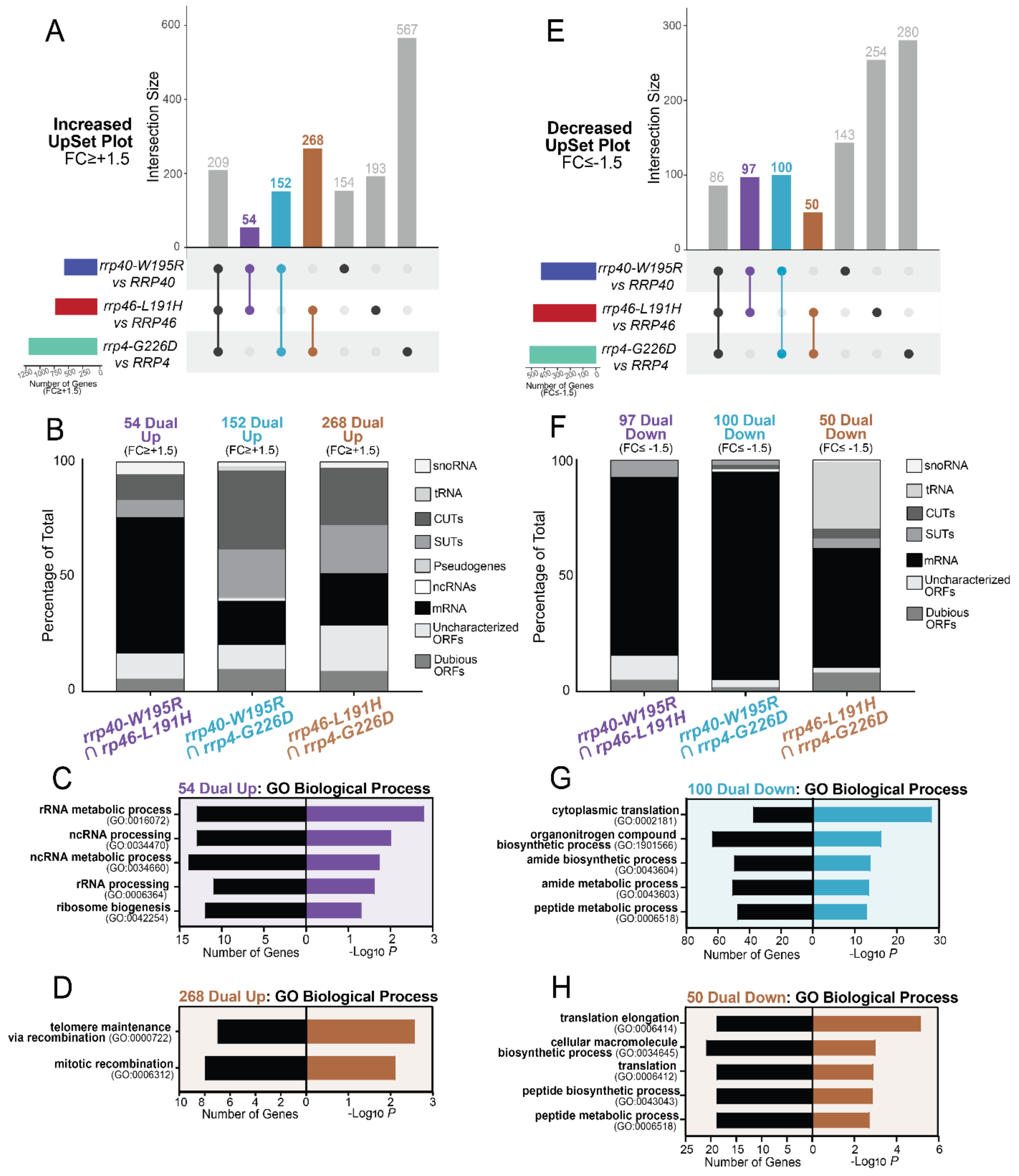

**Figure S4. UpSet Plots of differentially expressed transcripts in *rrp4-G226D, rrp40-W195R* and *rrp46-L191H* cells reveal targets shared differently between dual combinations of the three *rrp* mutants.** The UpSet plot of significantly increased (FC≥+1.5) (**A**) or decreased (FC≤-1.5) (**E**) transcripts. Stacked bar percentages of the RNA types that comprise the intersections of increased (Up) **(B)** or decreased transcripts (Down) (**F**) identified between the mutant pairs. (**C**) Gene ontology (GO) analysis: for biological processes of the 54 increased transcripts co-occurring in the *rrp40-W195R* and *rrp46-L191H* datasets; for biological processes of the 268 increased (**D**) or decreased (**H**) transcripts co-occuring in the *rrp4-G226D* and *rrp46-L191H* datasets; and for the 100 decreased transcripts co-occurring in the *rrp40-W195R* and *rrp4-G226D* datasets (**G**). In **C**, **D**, **G** and **H**- black bars represent the number of transcripts linked to each biological process category. Colored bars represent the -log of the associated *p*-value for each GO term. All GO analyses were performed on mRNAs and non-coding RNAs (tRNAs, snoRNAs, and snRNAs).

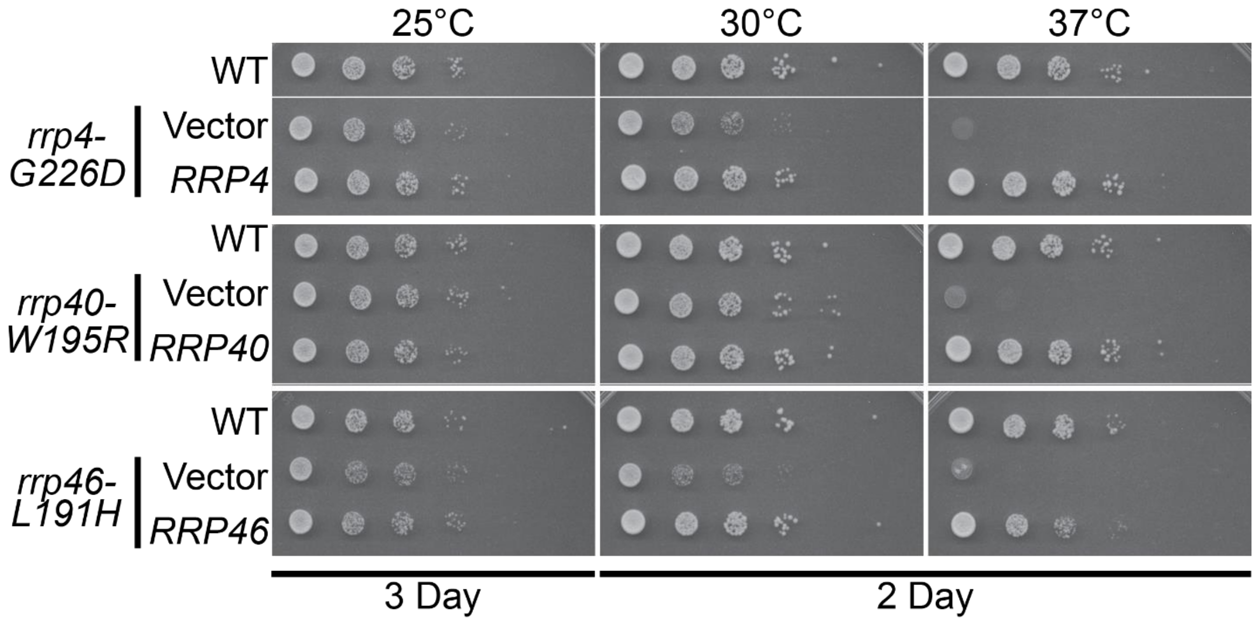


**Figure S5. Growth assays of CRISPR-engineered yeast strains expressing disease-associated RNA exosome mutants indicate severe growth defects for *rrp4-G226D, rrp40-W195R* and *rrp46-L191H* cells.** Cells were transformed with an empty vector or a plasmid expressing wild-type RNA exosome subunit before serially diluting and spotting onto solid YEPD media. Images shown are after three days of growth at 25°C and two days of growth at 30°C and 37°C. Parental wild-type cells (WT) were included as a control.

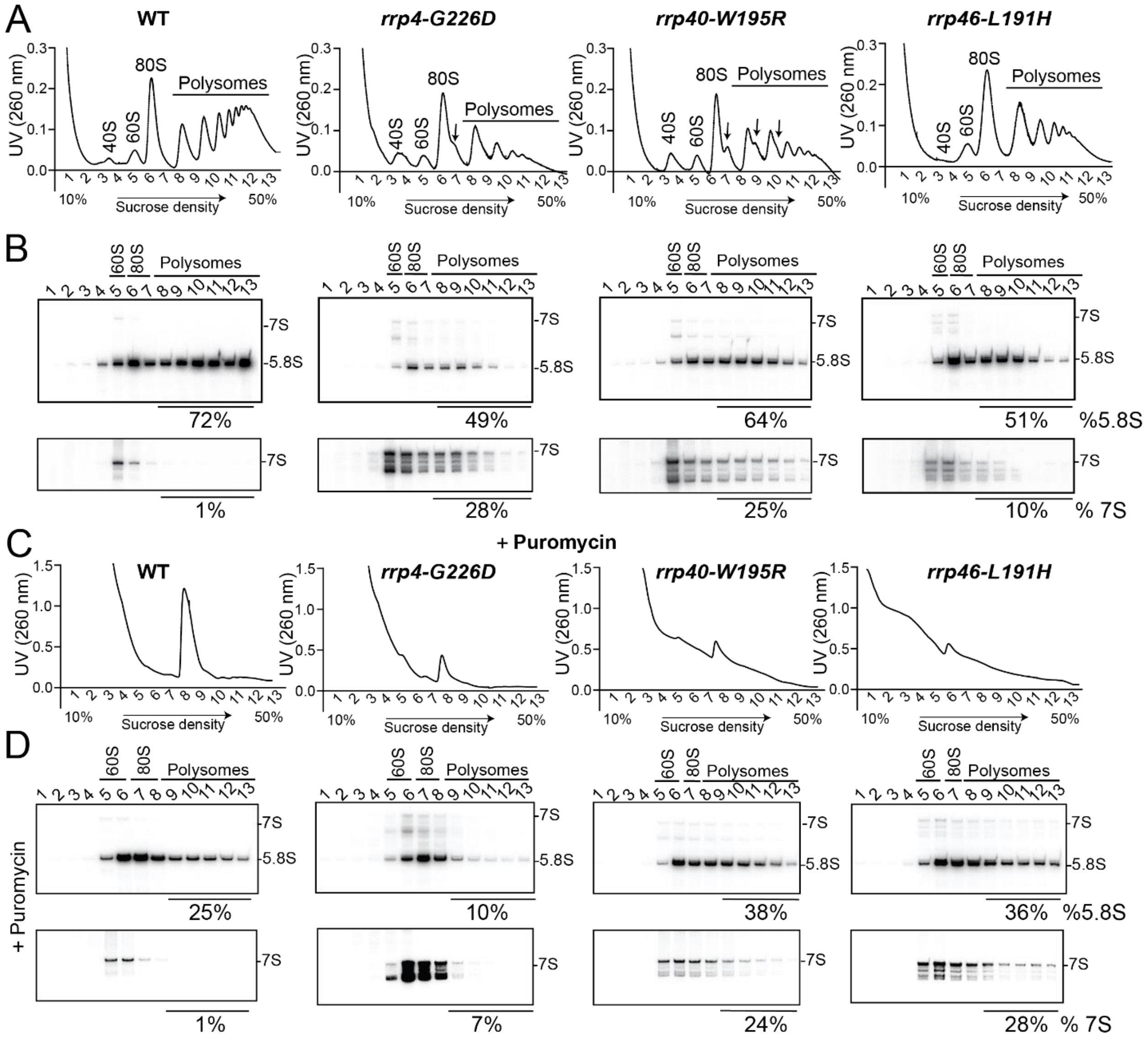


**Figure S6. Distinct translation defects in *rrp4-G226D, rrp40-W195R* and *rrp46-L191H* mutant cells at 37°C.** (**A**) Sucrose density gradients of wild-type, *rrp4-G226D*, *rrp40-W195R*, and *rrp46-L191H* cells that were grown 37°C are shown. Clarified cell extracts were resolved on a 10-50% sucrose gradient and scanned at 260 nm. Arrows indicate halfmers. (**B-C**) Northern blots of gradient fractions indicating distribution of 5.8S rRNA and 7S pre-rRNA. Samples in (**C**) were treated with 2.5 mM puromycin after lysis.

**Table S1. Disease-associated RNA exosome variants**

| Human Exosome Subunit Variant | Yeast Exosome Subunit Variant | Yeast Exosome Mutant Growth | | Yeast Exosome Protein Variant Level (% Rel. WT) | | Yeast 7S rRNA Accumulation | Reference |
|---|---|---|---|---|---|---|---|
| | | 30°C | 37°C | 30°C | 37°C | 37°C | |
| EXOSC2-G198D | Rrp4-G226D | Slow | V. Slow | 118% | 75% | Yes | (Sterrett et al. 2021) |
| EXOSC3-D132A | Rrp40-S87A | Normal | Normal | 99% | 68% | N.D. | (Fasken et al. 2017) |
| EXOSC3-W238R | Rrp40-W195R | Normal | Slow | 63% | 30% | Yes | (Fasken et al. 2017; Gillespie et al. 2017) |
| EXOSC5-T114I | Rrp46-Q86I | Normal | Normal | 58% | 52% | No | (Slavotinek et al. 2020) |
| EXOSC5-M148T | Rrp46-L127T | Normal | Normal | 90% | 60% | N.D. | (Slavotinek et al. 2020) |
| EXOSC5-L206H | Rrp46-L191H | Slow | Slow | 171% | 72% | Yes | (Slavotinek et al. 2020) |
| EXOSC9-L14P | Rrp45-I15P | Normal | Normal | N.D. | N.D. | N.D. | This Study |

**Table S2. *Saccharomyces cerevisiae* strains and plasmids**

| Strain/Plasmid | Description | Reference |
|---|---|---|
| *rrp4Δ* (yAV1103) | *MATα ura3Δ0 leu2Δ0 his3Δ1 lys2Δ0 rrp4Δ::NEO [RRP4, URA3]* | (Losh 2018) |
| *rrp40Δ* (yAV1107) | *MATa ura3Δ0 leu2Δ0 his3Δ1 rrp40Δ::NEO [RRP40, URA3]* | (Schaeffer et al. 2009) |
| *rrp45Δ* (yAV1410) | *MATα ura3Δ0 leu2Δ0 his3Δ1 rrp45Δ::NEO [RRP45, URA3]* | (Ahammed et al. 2025) |
| *rrp46Δ* (yAV1105) | *MATα ura3Δ0 leu2Δ0 his3Δ1 lys2Δ0 rrp46Δ::kanMX4 [RRP4, URA3]* | (Slavotinek et al. 2020) |
| BY4741 (ACY402) | *MATa ura3Δ0 leu2Δ0 his3Δ1 met15Δ0* | (Brachmann et al. 1998) |
| *rrp6Δ* (ACY1641) | *MATa ura3Δ0 leu2Δ0 his3Δ1 met15Δ0 rrp6Δ::kanMX4* | (Fasken et al. 2011) |
| *rrp4-G226D* (ACY3110) | *MATa ura3Δ0 leu2Δ0 his3Δ1 met15Δ0 rrp4-G226D* | This Study |
| *rrp40-W195R* (ACY3117) | *MATa ura3Δ0 leu2Δ0 his3Δ1 met15Δ0 rrp40-W195R* | This Study |
| *rrp46-L191H* (ACY3137) | *MATa ura3Δ0 leu2Δ0 his3Δ1 met15Δ0 rrp46-L191H* | This Study |
| pRS315 | *CEN6, LEU2, amp*$^R$ | |
| pRS316 | *CEN6, URA3, amp*$^R$ | (Sikorski and Hieter 1989) |
| pAC3656 | *RRP4-Native 3'UTR* in pRS315, *CEN6, LEU2, amp*$^R$ | (Sterrett et al. 2021) |
| pAC3659 | *rrp4-G226D-Native 3'UTR* in pRS315, *CEN6, LEU2, amp*$^R$ | (Sterrett et al. 2021) |

| | | |
|---|---|---|
| pAC3652 | *RRP40-Native 3'UTR* in pRS315*, CEN6, LEU2, amp$^R$* | (Sterrett et al. 2021) |
| pAC3654 | *rrp40-S87A-Native 3'UTR* in pRS315*, CEN6, LEU2, amp$^R$* | This Study |
| pAC3655 | *rrp40-W195R-Native 3'UTR* in pRS315*, CEN6, LEU2, amp$^R$* | (Sterrett et al. 2021) |
| pAV975/pAC3479 | *RRP45-Native 3'UTR* in pRS, *CEN6, LEU2, amp$^R$* | This Study |
| pAC3480 | *rrp45-I15P-Native 3'UTR* in pRS, *CEN6, LEU2, amp$^R$* | This Study |
| pAC3482 | *RRP46-Native 3'UTR* in pRS315, *CEN6, LEU2, amp$^R$* | (Slavotinek et al. 2020) |
| pAC3483 | *rrp46-Q86I-Native 3'UTR* in pRS315, *CEN6, LEU2, amp$^R$* | (Slavotinek et al. 2020) |
| pAC3484 | *rrp46-L191H-Native 3'UTR* in pRS315*, CEN6, LEU2, amp$^R$* | (Slavotinek et al. 2020) |
| pAC3534 | *rrp46-L127T-Native 3'UTR* in pRS315, *CEN6, LEU2, amp$^R$* | (Slavotinek et al. 2020) |
| pAC3752 | *RRP6* in pRS315*, CEN6, LEU2, amp$^R$* | This Study |
| pAC3846 | *TEF1p-Cas9-CYC1t-SNR52p* in pRS316, *CEN6, URA3, amp$^R$* | This Study |
| pAC3863 | *TEF1p-Cas9-CYC1t-SNR52p-RRP4_668.gRNA-SUP4t* in pRS316, *CEN6, URA3, amp$^R$* | This Study |
| pAC3861 | *TEF1p-Cas9-CYC1t-SNR52p-RRP40_583.gRNA-SUP4t* in pRS316, *CEN6, URA3, amp$^R$* | This Study |
| pAC4342 | *TEF1p-Cas9-CYC1t-SNR52p-RRP46_589.gRNA-SUP4t* in pRS316, *CEN6, URA3, amp$^R$* | This Study |
| pDB868 | PGK- Miscoding Firefly H245R in pYEplac195, *amp$^R$* | (Salas-Marco and Bedwell 2005) |
| pJD375 | ADH-0-Frame Control in pRS416*, amp$^R$* | (Harger and Dinman 2003) |
| pJD376 | ADH L-A (-1) Frameshift in pRS416*, amp$^R$* | (Harger and Dinman 2003) |
| pJD377 | ADH Ty1 (+1) Frameshift in pRS416*, amp$^R$* | (Harger and Dinman 2003) |
| pRaugFuug | ADH/GPD start in pRS416*, amp$^R$* | (Cheung et al. 2007) |
| pRaugFaug | ADH/GPD start in pRS416*, amp$^R$* | (Cheung et al. 2007) |

**Table S3. List of oligonucleotides used in this study**

| Description | Sequence (5'-3') | Name |
|---|---|---|
| *rrp45-I15P* F | CCGCATCCGAGTCAAAATTTCCCTTAGAAGCACTGAGACAG AATTATAGG | AC8801 |
| *rrp45-I15P* R | CCTATAATTCTGTCTCAGTGCTTCTAAGGGAAATTTTGACT CGGATGCGG | AC8802 |
| *TEF1p* F | ATATGAGCTCATAGCTTCAAAATGTTTCTACTCC | AC8410 |
| *TEF1p* R | ATATACTAGTAAAACTTAGATTAGATTGCTATGC | AC8411 |

| | | |
|---|---|---|
| *Cas9* F | ATATACTAGTAAAATTTTGGGCCCAAAAAAATGGACAAGAAGTACTCCATTGGGCTCGATATCGG | AC6802 |
| *CYC1t* R | ATATGGTACCAAAATTTTACCGGTGGCCGCAAATTAAAGCCTTCGAGCGTCCCAAAACCTTCTCAAGCAAGG | AC6803 |
| *SNR52p* F | ATATACCGGTGGCACCCCAGGCTTTACACTTTATGCTTCCGG | AC6804 |
| *SNR52p* R | ATATGGTACCTTTTAAAAGCATGCGATCATTTATCTTTCACTGCGGAGAAGTTTCG | AC6805 |
| *RRP4_668-gRNA* F | ATATGCATGCACCATTGACTCCGAGAACTAGTTTTAGAGCTAGAAATAGC | AC8407 |
| *RRP40_583-gRNA* F | ATATGCATGCTGGTCTCAATGGGAAGATCTGTTTTAGAGCTAGAAATAGC | AC8402 |
| *RRP46_589-gRNA* F | ATATGCATGCATTGTTCAGTTTACTGGAGCGTTTTAGAGCTAGAAATAGC | AC9888 |
| *SUP4t-crRNA* R | ATATGGTACCAGACATAAAAAACAAAAAAAGCACCACC | AC6809 |
| *rrp4-G226D* _HDR F | GAACCATACTCATAATTTGCCCGGGAACATAACAGTAGTTCTCGATGTCAATGGTTACATATGGTTAAGGAAAACATCTCAGATGGAC | AC8408 |
| *rrp4-G226D* _HDR R | GTCCATCTGAGATGTTTTCCTTAACCATATGTAACCATTGACATCGAGAACTACTGTTATGTTCCCGGGCAAATTATGAGTATGGTTC | AC8409 |
| *rrp40-W195R* _HDR F | TACCAAGTTTGAAGTCGCCATTGGTCTCAATGGGAAGATCAGAGTTAAGTGCGAAGAATTATCTAACACTTTAGCTTGTTATAGAACC | AC8404 |
| *rrp40-W195R* _HDR R | GGTTCTATAACAAGCTAAAGTGTTAGATAATTCTTCGCACTTAACTCTGATCTTCCCATTGAGACCAATGGCGACTTCAAACTTGGTA | AC8405 |
| *rrp46-L191H* _HDR F | TAGACAGCAATGGTGATTTTAATGAAGATCAACACTTCAGTTTACTAGAGCTAGGTGAGCAAAAGTGTCAAGAACTTGTCACAAATAT | AC9889 |
| *rrp46-L191H* _HDR R | ATATTTGTGACAAGTTCTTGACACTTTTGCTCACCTAGCTCTAGTAAACTGAAGTGTTGATCTTCATTAAAATCACCATTGCTGTCTA | AC9890 |
| Probe 18S | CATGGCTTAATCTTTGAGAC | HG243 |
| Probe 25S | GCCCGTTCCCTTGGCTGTG | HG244 |
| Probe 5.8S | CTGCGTTCTTGATCGATGCG | HG247 |
| Probe 5S | CTACTCGGTCAGGCTC | HG246 |
| Probe ITS2 (within ITS2) | GGCCAGCAATTTCAAGTTA | HG242 |
| Probe ITS2a (binds at the end of 5.8S and beginning of ITS2) | TGAGAAGGAAATGACGCT | HG1066 |
| Probe ITS1a (between A2 and A3) | TGTTACCTCTGGGCCC | HG245 |
| Probe ITS1b (between D and A2) | GCTCTCATGCTCTTGCC | HG240 |

### References


Ahammed KS, Fasken MB, Corbett AH, van Hoof A. 2025. Humanized Saccharomyces cerevisiae provides a facile and effective tool to identify damaging human variants that cause exosomopathies. *G3 (Bethesda)*.

Brachmann CB, Davies A, Cost GJ, Caputo E, Li J, Hieter P, Boeke JD. 1998. Designer deletion strains derived from Saccharomyces cerevisiae S288C: a useful set of strains and plasmids for PCR-mediated gene disruption and other applications. *Yeast* **14**: 115-132.

Cheung YN, Maag D, Mitchell SF, Fekete CA, Algire MA, Takacs JE, Shirokikh N, Pestova T, Lorsch JR, Hinnebusch AG. 2007. Dissociation of eIF1 from the 40S ribosomal subunit is a key step in start codon selection in vivo. *Genes Dev* **21**: 1217-1230.

Fasken MB, Leung SW, Banerjee A, Kodani MO, Chavez R, Bowman EA, Purohit MK, Rubinson ME, Rubinson EH, Corbett AH. 2011. Air1 zinc knuckles 4 and 5 and a conserved IWRXY motif are critical for the function and integrity of the Trf4/5-Air1/2-Mtr4 polyadenylation (TRAMP) RNA quality control complex. *J Biol Chem* **286**: 37429-37445.

Fasken MB, Losh JS, Leung SW, Brutus S, Avin B, Vaught JC, Potter-Birriel J, Craig T, Conn GL, Mills-Lujan K et al. 2017. Insight into the RNA Exosome Complex Through Modeling Pontocerebellar Hypoplasia Type 1b Disease Mutations in Yeast. *Genetics* **205**: 221-237.

Gillespie A, Gabunilas J, Jen JC, Chanfreau GF. 2017. Mutations of EXOSC3/Rrp40p associated with neurological diseases impact ribosomal RNA processing functions of the exosome in S. cerevisiae. *Rna* **23**: 466-472.

Harger JW, Dinman JD. 2003. An in vivo dual-luciferase assay system for studying translational recoding in the yeast Saccharomyces cerevisiae. *Rna* **9**: 1019-1024.

Losh J. 2018. Identifying Subunit Organization and Function of the Nuclear RNA Exosome Machinery. In *UTHealth Graduate School of Biomedical Sciences Dissertations and Theses*. The University of Texas MD Anderson Cancer Center Open Access.

Salas-Marco J, Bedwell DM. 2005. Discrimination between defects in elongation fidelity and termination efficiency provides mechanistic insights into translational readthrough. *J Mol Biol* **348**: 801-815.

Schaeffer D, Tsanova B, Barbas A, Reis FP, Dastidar EG, Sanchez-Rotunno M, Arraiano CM, van Hoof A. 2009. The exosome contains domains with specific endoribonuclease, exoribonuclease and cytoplasmic mRNA decay activities. *Nat Struct Mol Biol* **16**: 56-62.

Sikorski RS, Hieter P. 1989. A system of shuttle vectors and yeast host strains designed for efficient manipulation of DNA in Saccharomyces cerevisiae. *Genetics* **122**: 19-27.

Slavotinek A, Misceo D, Htun S, Mathisen L, Frengen E, Foreman M, Hurtig JE, Enyenihi L, Sterrett MC, Leung SW et al. 2020. Biallelic variants in the RNA exosome gene EXOSC5 are associated with developmental delays, short stature, cerebellar hypoplasia and motor weakness. *Human molecular genetics* **29**: 2218-2239.

Sterrett MC, Enyenihi L, Leung SW, Hess L, Strassler SE, Farchi D, Lee RS, Withers ES, Kremsky I, Baker RE et al. 2021. A budding yeast model for human disease mutations in the EXOSC2 cap subunit of the RNA exosome complex. *RNA* **27**: 1046-1067.
